# Complement-associated loss of CA2 inhibitory synapses in the demyelinated hippocampus impairs memory

**DOI:** 10.1101/2021.01.15.426022

**Authors:** Valeria Ramaglia, Mohit Dubey, M. Alfonso Malpede, Naomi Petersen, Sharon I. de Vries, Dennis S.W. Lee, Geert J. Schenk, Stefan M. Gold, Inge Huitinga, Jennifer L. Gommerman, Jeroen J.G. Geurts, Maarten H.P. Kole

## Abstract

The complement system is implicated in synapse loss in the MS hippocampus, but the functional consequences of synapse loss remain poorly understood. Here, in post-mortem MS hippocampi with demyelination we find that deposits of the complement component C1q are enriched in the CA2 subfield, are linked to loss of inhibitory synapses and are significantly higher in MS patients with cognitive impairments compared to those with preserved cognitive functions. Using the cuprizone mouse model of demyelination, we corroborated that C1q deposits are highest within the demyelinated dorsal hippocampal CA2 pyramidal layer, and co-localized with inhibitory synapses engulfed by microglia/macrophages. In agreement with the loss of inhibitory perisomatic synapses, we further found that Schaffer collateral feedforward inhibition but not excitation was impaired in CA2 pyramidal neurons and accompanied by a reduced spike output. Ultimately, we show that these electrophysiological changes were associated with an impaired encoding of social memories. Together, our findings identify CA2 as a critical circuit in demyelinated intrahippocampal lesions and memory dysfunctions in MS.

## Introduction

Multiple sclerosis (MS) is an autoimmune demyelinating and neurodegenerative disease of the central nervous system (CNS). Patients living with MS experience major cognitive disabilities including memory impairment, attention deficits and slowed sensory processing speed [1, 2], which occurs from the early stages of the disease [3]. Recent emerging insights have drawn particular attention to MS-related deficits in social cognition and facial emotion recognition as affected cognitive domains in MS, which can occur early in disease even in the absence of other cognitive problems [4] and may have distinct neuropathological substrates [5, 6]. There is substantial evidence that the hippocampus is critical for the consolidation and recollection of episodic memories, the temporal organization of events and mapping of social space [7, 8]. Recent magnetic resonance imaging (MRI) studies have shown that structural and functional disconnections of the hippocampus from several brain networks can explain some of the clinical deficits experienced by MS patients including impaired memory and learning [9, 10] as well as depressive symptoms [11, 12]. In addition, post-mortem studies reported that the hippocampus of MS patients often shows extensive demyelination [13, 14].

The molecular and cellular basis of the MS-related hippocampal damage is, however, not fully understood. One leading hypothesis based on experimental [15, 16] and post-mortem studies [16, 17] indicates that the disconnection of temporal lobe networks in MS may be due to the loss of synapses via a “pruning” process. Over the past decade, several studies have identified proteins of the complement system as key components of the pruning process in development and learning [18-23]. The complement system is traditionally known as a major arm of the innate immune system, required for optimal defense against pathogens and for the disposal of dead and dying cells [24]. The recently discovered role for complement in developmental synaptic pruning has been extensively investigated in the retinogeniculate system, where exuberant and overlapping synaptic connections are progressively segregated into eye-specific projections [25]. In this system, supernumerary synapses are targeted by the complement component 1q (C1q), opsonized by C3, and phagocytosed by microglia via complement receptor 3 (CR3) [18, 19]. In the rodent brain C1q has also been shown to play a role in shaping synaptic circuits in memory formation during adulthood [26], in ageing [27] and in neurodegeneration [22]. A more recent report showed that synaptic material is tagged by C3 (but not by C1q) and is engulfed by microglia in the retinogeniculate system of models of demyelination and in the visual thalamus of MS patients [16]. Our team previously showed that in the MS hippocampus C1q and C3d are deposited particularly in the CA2/3 region at synapses that localize within microglial processes and lysosomes, supporting a role for microglia in the elimination and degradation of synapses [28]. However, the nature of the targeted circuits, the mechanisms of synapse elimination and the functional consequences of synapse loss in the MS hippocampus are unknown.

Here, we first evaluated the extent of C1q depositions in CA2 versus CA3 regions of myelinated and demyelinated MS hippocampi, and related it to synaptic changes, MRI measures of brain atrophy and the cognitive status of the patient. To further investigate the significance of C1q deposition and synaptic changes observed in the MS hippocampus, we studied the extent and localization of C1q/C3 proteins in relation to synapses in the cuprizone model. Electrophysiological recordings were acquired, and social memory performance was tested to evaluate the consequences of subfield-specific synaptic changes in cuprizone-fed mice. Together, our findings identify the CA2 region of the hippocampus as a subfield that is highly susceptible to complement deposition and synaptic reorganization of inhibitory circuit in MS. These changes may play a critical role in altering hippocampal information flow underlying cognitive deficits in the social domain.

## Materials & Methods

### Human studies

#### Post-mortem hippocampal tissue collection

Post-mortem hippocampi of 55 MS donors and 5 non-neurological control (NNC) donors were obtained from the Netherlands Brain Bank (NBB; Amsterdam, the Netherlands). NBB autopsy procedures were approved by the Independent Review Board of the Amsterdam UMC, registered with US office of Human Research Protections. Written informed consent was obtained by the NBB for brain autopsy and for the use of material and clinical data for research purposes, in compliance with institutional and national ethical guidelines. Brains were removed according to a rapid (post-mortem delay (PMDS) of 6.7 ± 3.2 hours, mean ± SD) autopsy protocol. Specimens were fixed in 10% buffered formalin and processed for embedding in paraffin. Paraffin-embedded hippocampi of all donors were used for the pathological study. Of these, hippocampi of 14 MS donors were selected for pathological-MRI correlation studies and hippocampi of 18 MS donors were selected for pathological-clinical correlation studies. Coronally cut hippocampi were selected to ensure accurate and systematic scoring of demyelination, C1q deposition and synaptic changes within the anatomical subfields of the hippocampus. MS cases and controls were matched for age, all numbers represent mean ± SD (MS myelinated [MS-M] donors: 65.2 ± 12.0 years; MS demyelinated [MS-DM] donors: 63.5 ± 15.3 years; NNC donors: 64.4 ± 15.4 years; one-way ANOVA *P* = 0.94) and PMD (MS-M donors: 7.4 ± 3.8 hours; MS-D donors: 5.8 ± 2.3 hours; NNC donors: 6.9 ± 1.7 hours; one-way ANOVA *P* = 0.15). MS paraffin-embedded hippocampi used for immunostaining were from 29 donors with primary (PP) or secondary progressive (SP) disease and 26 donors with progressive disease of undetermined type (PP/SP). In this study, PP, SP, and PP/SP donors were pooled and referred to as progressive MS. Detailed clinicopathological data of all donors are provided in **Supplementary Table 1**.

#### Magnetic resonance imaging (MRI)

The MRI protocol comprises both whole-brain in-situ MRI, and MRI of 10-mm thick coronal brain slices, which are cut at autopsy. A detailed description was previously published [29]. MR imaging was performed using 1.5T Siemens Sonata and Avanto MRI scanners, depending on the availability at the time of autopsy, as described previously [30]. Briefly, the *in-situ* image acquisition protocol for volumetry of the hippocampus included a sagittal 3DT1-weighted imaging sequence (TR = 2700 ms, TE = 5.1 ms, TI = 950 ms, FA = 8, voxel size = 1.2 × 1.2 × 1.3 mm) and a sagittal 3D-FLAIR sequence (TR = 6500 ms, TE = 355 ms, TI = 2200 ms, voxel size = 1.2 × 1.2 × 1.3 mm). The 3DT1 images were used to measure whole hippocampus volumes corresponding with the hemisphere of the tissue samples extracted for neuropathological assessment using the FreeSurfer image analysis suite version 5.3, which is documented and freely available for download online (http://surfer.nmr.mgh.harvard.edu/).

#### Evaluation of cognitive function. Evaluation of cognitive function

Inspecting the clinical data of all cases included in the MS post-mortem collection of the Netherlands Brain Bank (http://www.brainbank.nl/), we identified MS cases for which neuropsychological information was available. Using clinical chart information on cognitive status has proven successful in post-mortem research before [13]. The demographic and clinical data of the selected CP and CI cases are summarized in **Supplementary Table 1**. By excluding any cases 1) without detailed information on cognition, 2) with a neuropsychological history (e.g. depression, character changes) and 3) that had any other non-MS pathology (e.g. vascular pathology), we were able to select high quality post-mortem material from cognitively preserved (CP; *n* = 7) and cognitively impaired (CI; *n* = 7) MS patients. All included CI patients had memory problems that were often accompanied by disturbed linguistic capabilities.

#### Hippocampal Lesion Classification

Hippocampal tissue sections were stained for proteolipid protein (PLP) and for the anti–human leukocyte antigen (HLA_DP-DQ-DR_). Because the distribution of HLA_DP-DQ-DR_–immunopositive microglial cells did not segregate with lesional areas, samples were scored for the presence of lesions according to their anatomical location and not lesion activity. Only intrahippocampal lesions were scored.

#### Immunohistochemistry

For the immunohistochemistry, endogenous peroxidase activity was blocked by incubating the slides in methanol with 0.3% H_2_O_2_ for 20 minutes at room temperature (RT). Sections were washed in 1× PBS (9 minutes) and put in a microwave on “High” power settings for 20 minutes in 10 mM Tris/1 mM Ethylenediaminetetraacetic acid (EDTA) buffer pH.9 **(Supplementary Table 2)**. Sections were rinsed in 1× PBS, outlined with a hydrophobic pen, washed in 1× PBS and PBST (3 minutes). The sections were then blocked with normal goat serum in PBST (1:1) for 30 minutes at RT before being incubated with the relevant primary antibody **(Supplementary Table 2)** diluted in Normal Antibody Diluent (Immunologic, Duiven, The Netherlands) for 1 hour at RT and then overnight at 4°C. The next day, slides were rinsed in PBST (9 minutes) and incubated with Post Antibody Blocking BrightVision Solution 1 (diluted 1:1 in PBST, ImmunoLogic) for 15 minutes at RT. They are then washed in 1× PBS and incubated with BrightVision Poly-HRP-Anti Ms/Rb/Rt IgG Biotin-free Solution 2 (diluted 1:1 in PBST, ImmunoLogic) for 30 minutes at RT. The immunostaining was visualised using 3,3’-Diaminobenzidine (DAB, Sigma-Aldrich) for 4 minutes at RT. The sections were counterstained with hematoxylin. Sections were then dehydrated in a series (50%, 70%, 100%, 100%) of ethanol and xylene (3 minutes). The slides were mounted using Entellan medium. All stained images were scanned using an Axio Imager Z1, Zeiss microscope connected to a digital camera (AxioCam 506 mono, Zeiss) and imaged with Zen pro 2.0 imaging software (Zeiss).

#### Immunofluorescence staining of synapses

For the fluorescent immunostaining of pre- and postsynaptic elements, sections were pretreated with microwave antigen retrieval as described above. Primary antibodies against the presynaptic elements vesicular glutamate transporter 1 (vGLUT1) or vesicular GABA transporter (vGAT) (see **Supplementary Table 2**) were diluted in normal antibody diluent (Immunologic, Duiven, the Netherlands) and incubated for 3 hours at RT followed by overnight incubation at 4°C. The next day, sections were washed in PBS and incubated in primary antibodies against the postsynaptic elements postsynaptic domain 95 (PSD95) or Gephyrin (see **Supplementary Table 2**) diluted in normal antibody diluent for 4 hours at RT followed by 2 overnights incubation at 4°C. Two days later, sections were washed in PBS and incubated in polyclonal IgG donkey anti-guinea pig Alexa488-conjugated (Jackson, A-S155) and the polyclonal IgG donkey anti-rabbit Alexa546-conjugated (Invitrogen, A-S154) secondary antibodies or the polyclonal IgG donkey anti-chicken Alexa488-conjugated (Jackson, A-S153) and the polyclonal IgG donkey anti-mouse Alexa546-conjugated (Molecular probes, A-S032) secondary antibodies diluted 1:200 in PBS supplemented with 3% donkey serum with 0.1% triton for 3 hours at RT. After washing in PBS, sections were incubated with 40,6-diamidino-2-phenylindole (DAPI; Vector Laboratories) to visualize the nuclei, incubated in Sudan Black B for 5 minutes at RT. After washing in 70% ethanol and sqH_2_O, the slides were air dried and mounted in aqueous mounting medium. Using the appropriate filters, the immunofluorescence signal was visualized with an Axio Imager Z1, Zeiss microscope connected to a digital camera (AxioCam 506 mono, Zeiss) and imaged with Zen pro 2.0 imaging software (Zeiss). To control for antibodies specificity, tissue sections were stained according to the IF or IHC protocols described above except for the primary antibody incubation step, which was omitted.

#### Quantification of immunohistochemistry

Formalin-fixed paraffin embedded tissue blocks were cut into 7µm sections on a microtome (ThermoScientific HM 325), mounted onto Superfrost Plus glass slides and dried overnight at 37°C. Sections were deparaffinized in xylene (2 × 5 minutes), rehydrated through a series (100%, 70%, 50%) of ethanol and sqH_2_O (3 minutes). For the CA2 and CA3 subfields 3 randomly selected nonoverlapping digital images were captured for quantification. Therefore, for each immunostaining, a total of 30 images (3 images × 2 subfields ×5 donors) of NNCs, 186 images (3 images × 2 subfields × 31 donors) of MS-M hippocampi, and 144 images (3 images × 2 subfields × 24 donors) of MS-DM hippocampi were captured at x20 magnification and analysed. Quantitative analysis of immunostaining was performed on the region of interest (ROI) using the measurement function of ImageJ 1.15s (National Institutes of Health). Briefly, the RGB images were separated into single color channels using the color deconvolution plugin in Image J. The single-color channel for each staining was subjected to thresholding to create a mask that captures the specific staining. The threshold was saved and applied to all images in the same staining group. The area fraction measurement was applied to each ROI to quantify the percentage of thresholded staining. The amount of staining is expressed as percentage of immunoreactive area over the total area assessed.

### Animal studies

All animal experiments were performed in compliance with the European Communities Council Directive 2010/63/EU effective from 1 January 2013. The experiments were evaluated by the KNAW Animal Ethics Committee (DEC) and Central Authority for Scientific Procedures on Animals (CCD, license AVD8010020172426). The specific experimental designs were evaluated and monitored by the Animal Welfare Body (IvD, protocols NIN18.21.01, NIN19.21.06 and NIN19.21.07). Male C57BL/6 mice (Janvier Labs, Saint-Berthevin Cedex, France) were kept on a 12:12 h light-dark cycle (lights on at 07.00 am, lights off at 19.00 pm) with ad libitum food and water. Demyelination was induced by cuprizone feeding [31]. From the age of 5–6 weeks and a bodyweight >20 grams (on average 21.6 g, range: 20.5 – 22.8 g), mice were fed with 0.2% cuprizone supplemented to the powder food, freshly prepared every second day for a period between 2 and 9 weeks as indicated in the text. The associated weight loss with cuprizone treatment was assessed every second day and monitored in consultation with the IvD.

#### Behavioral tests

The *five-trial social memory test* was based on the design from Hitti & Siegelbaum [32]. All mice (*n* = 11 control and 11 cuprizone treated mice, 0.2% for 7 weeks) were maintained group-housed (3–4 mice/cage) before the test and the sequence of testing was determined randomly. Social memory tests were performed between 08.00 am and 15.00 pm. For the test the subject mouse was transferred to the experimental room and allowed to familiarize with the cage for 15 minutes with the lid closed. After 15 minutes, the lid was removed, and the webcam recording started (∼30 Hz frame rate). At this point, the subject mouse was exposed to a novel male mouse for the duration of 1 minute (trial 1). The novel mouse was removed for 10 minutes. Subsequently, the same procedure was repeated three more times (i.e. subject mouse exposed to the familiar mouse, trials 2, 3 and 4). In trial 5, an unfamiliar mouse was introduced to measure dishabituation. The behaviors of the subject mouse were analysed off-line. The behavioural scoring included the duration of anogenital sniffing, approaching behavior, social interaction, aggressive interaction or no interaction. The occurrence and durations of these distinct behaviors were measured by two different researchers, both blinded to the animal identities, until the data were analysed and plotted.

For automated *discrimination learning* experiments (Sylics Bioinformatics, Amsterdam, The Netherlands) we used PhenoTyper cages (model 3000, Noldus Information Technology). The system is an automated home cage in which behavior is tracked by a video. The cage is equipped with a drinking bottle and a triangular-shaped shelter with two entrances in one corner. In the opposite corner, an aluminum tube of an automated food reward dispenser protruded into the cage. Mice (*n* = 9 control and 13 cuprizone-treated mice, 0.2% for 6 weeks) had *ad libitum* access to drinking water but needed to engage for food reward in the Cognition Wall discrimination test. The wall contained three entrances and when they passed through the left entrance, they automatically obtained a food pellet (Dustless Precision Pellets, 14 mg, Bio-Serve). The rate at which a mouse gains a relative preference for the rewarded entrance is used as a measure of discrimination learning. Mice were single housed for one week to accommodate to the 16 hours periods in which they were housed in the PhenoTyper cages and experiment started 3 hours before lights-off (16.00 pm). C57BL/6J mice require typically around 100 food rewards/per day to maintain body weight. We analyzed the total number of entries needed to reach a criterion of 70% to 90% correct, computed as a moving window over the last 30 entries to assess learning in the task. Since this performance measurement uses the fraction of correct over incorrect entries in the last 30 entries rather than the total number of entries or latency to reach criterion, this measurement is not likely to be influenced by general differences in activity between genotypes or groups. Hence, mice cannot achieve the learning criterion by only showing increased motor activity and making more entries.

#### Hippocampus slice preparation and electrophysiological recordings

Mice received a terminal dose of Nembutal (5 mg kg^−1^) and were transcardially perfused with ice-cold artificial cerebrospinal fluid (ACSF) consisting of (in mM): 87.0 NaCl, 25.0 NaHCO_3,_ 2.5 KCl, 25.0 NaH_2_PO_4_, 75.0 sucrose, 25.0 glucose, 0.5 CaCl_2_ and 7.0 MgCl_2_ (oxygenated with 5% CO_2_–95% O_2_, pH 7.4). And after 20 minutes replaced for storage solutions containing 125 NaCl, 3 KCl, 25 glucose, 25 NaHCO_3_, 1.25 Na_2_H_2_PO_4_, 1 CaCl_2_, 6 MgCl_2_, 1 kynurenic acid (95% O_2_ and 5% CO_2_, pH 7.4). After decapitation, the brain was quickly removed from the skull and the hippocampus was isolated from the inside of the cortical mantle in an ice-cold (0 to +4°C) dissecting solution. The isolated hippocampus was placed in the groove of an agar block with the anterior part facing upward. Transverse hippocampal sections (400 µm) were cut starting at the dorsal site of the hippocampus using a Vibratome (1200VT, Leica Microsystems). Slices were allowed a recovery period of 30 min at 35°C and were subsequently stored at room temperature. For patch-clamp recordings, slices were transferred to an upright microscope (BX51WI, Olympus Nederland) equipped with oblique illumination optics (WI-OBCD; numerical aperture, 0.8). CA2 pyramidal cells located deep in the slice were visualized using 40× water-immersion objectives (Olympus) and oblique LED illumination optics (850 nm) based on the curvature of the pyramidal layer, the typical large soma and triangle shape. Some neurons showed a proximal bifurcation in the main apical dendrite. The microscope bath was perfused with oxygenated (95% O_2_, 5% CO_2_) ACSF consisting of the following (in mM): 125 NaCl, 3 KCl, 25 D-glucose, 25 NaHCO_3_, 1.25 Na_2_H_2_PO_4_, 2 CaCl_2_, and 1 MgCl_2_.

Patch pipettes were pulled from borosilicate glass (Harvard Apparatus, Edenbridge, Kent, UK) to an open tip of 3–6 MΩ resistance. For all current-clamp recordings the intracellular solution contained (in mM): 130 K-Gluconate, 10 KCl, 4 Mg-ATP, 0.3 Na_2_-GTP, 10 HEPES and 10 Na_2_-phosphocreatine (pH 7.25 adjusted with KOH, 280 mOsmol kg^−1^). The liquid junction potential difference of –13.5 mV was corrected in all recordings. All voltage recordings were analogue low-pass filtered at 10 kHz (Bessel), recorded using DAGAN BVC 700 amplifiers and digitally sampled at 100 kHz using an ITC-18 A-D converter (HEKA Elektronik Dr. Schulze GmbH, Germany). Bridge-balance and pipette or stray capacitances were fully compensated based on small current injections leading to minimal voltage errors. All data acquisition and analyses were performed with Axograph X (v.1.7.0, NSW, Australia, https://www.axograph.com/). The recording temperature was 34 ± 1°C. Only cells with a stable bridge-balance (< 25 MΩ) and resting membrane potential throughout the recording session were included in the analyses.

#### Morphological analysis and pyramidal cell identification

For single cell biocytin-labelling, recorded neurons were filled with 5 mg ml^−1^ biocytin for at least 30 minutes. Streptavidin biotin-binding protein (Streptavidin Alexa 488, 1:500, Invitrogen) was diluted in 5% BSA with 5% NGS and 0.3% Triton-X overnight at 4°C. To identify the CA2 region, primary antibody rabbit anti-PCP4 (1:250 Sigma Aldrich, HPA005792) or mouse anti-RGS14 (1:500, Neuromab) were added to an overnight incubation mix. Secondary antibodies were Alexa 633 goat anti-rabbit (1:500; Invitrogen) or Alexa 633 goat anti-mouse (1:500; Invitrogen). Brain slices were mounted on glass slides and cover slipped with Vectashield H1000 fluorescent mounting medium (Vector Laboratories, Peterborough, UK) and sealed. Sections were imaged using a confocal laser-scanning microscope (SP8, DM6000 CFS; acquisition software, Leica Application Suite AF v3.2.1.9702) with 40× oil-immersion objectives and 1× digital zoom with step size of 0.5 µm. Alexa 488 and Alexa 633 were imaged using 488 and 633 excitation wavelengths, respectively. Confocal *z*-stacks were imported into Neurolucida 360 software (v2020, MBF Bioscience) for reconstruction using the interactive user-guided trace with the Directional Kernels method. Axon and basal and apical dendrite segments were analyzed using Neurolucida Explorer (MBF Bioscience).

#### Immunohistochemistry and synapse staining

Mice were anaesthetised with Nembutal (5 mg kg^−1^), the brain rapidly removed and immersion-fixed with 4% PFA overnight at room temperature. The fixed brains were briefly rinsed in PBS (Phosphate Buffer Solution) before sunken in 30% sucrose/PBS solution at 4°C, frozen with dry ice and stored at –80°C. One day before the experiment the frozen brains were moved to –20°C and stored overnight. The day of the experiment 14 µm sagittal or coronal sections were produced with a freezing-sliding microtome and stored in PBS at 4°C. Free floating sections were permeabilized at RT with 10% normal goat serum in 0.3% Triton X-100 in PBS for 2 h, followed by primary antibody incubation overnight at 4°C. Primary antibodies used, dilution and sources are provided in **Supplementary Table 2**. After rinsing 3× in PBS for 15 min, sections were incubated with secondary antibodies (1:500) in PBS with 3% goat serum for 2 h at room temperature. After rinsing 3× in PBS for 15 min, sections were mounted on glass slides, using vectashield containing DAPI (Vector labs H-1000). Fluorescence signals were imaged with a Leica TCS SP5 II (DMI6000 CFS; acquisition software Leica Application Suite AF v. 2.6.3.8173) or SP8 confocal laser-scanning microscope (DM6000 CFS; acquisition software, Leica Application Suite AF v3.2.1.9702, Leica Microsystems GmbH). Confocal images used for the intensity analysis were acquired at 1,096 × 1,096 pixels (2.0 or 3.0 μm *z*-step) using a 10× objective. Density of puncta were acquired at 1,096 × 1,096 pixels (2.0 or 3.0 μm *z*-step) using a 40× or 63× oil-immersion objectives (0.75–1.0 digital zoom). To avoid bleed through between emission wavelengths, automated sequential acquisition of multiple channels was used, and images saved as uncompressed LIF format.

#### Electron Microscopy

Tissue for electron microscopy was obtained from adult mice that were transcardially perfused and postfixed with freshly prepared 2% PFA and 2.5% glutaraldehyde in a 0.1M phosphate buffer (PB) pH 7.4. All steps were done at room temperature, unless stated otherwise. After subsequent washes in PB, tissues were cryo-protected through a gradient of 10%, 20% and 30% sucrose in PB and frozen on aluminium boats on dry ice. Coronal sections of 40 μm containing the hippocampus were obtained using a freezing microtome. Frozen coronal sections of the hippocampus were washed in PB, slices were blocked 2 hours with 5% normal goat serum in PB and incubated overnight with rabbit-α-C1q antibody (1:1000 in blocking solution) while shaking. Slices were washed in PB, incubated for 2 hours with a horseradish peroxidase coupled rabbit secondary antibody, washed in PB, pre-incubated for 20 minutes with 0.05% 3,3′-diaminobenzidine (DAB) in PB and incubated for 5’ with DAB and 0.03% H_2_O_2_ for visualization. The DAB reaction product was then intensified using the gold-substituted silver peroxidase method as previously described [33]. Briefly, slices were rinsed in 2% sodium acetate buffer and incubated for 2 hours in 10% sodium thioglycolate on a shaker. After multiple washes with the acetate buffer, slices were incubated for 6 minutes with silver solution, consisting of 2.5% sodium carbonate, 0.1% ammonium nitrate, 0.1% silver nitrate, 0.5% tungstenosilic acid and 0.07% formalin. Following washes with acetate buffer, slices were incubated with 0.05% gold chloride for 20 min, rinsed with acetate buffer and incubated with 3% sodium thiosulfate for 5 min. After rinsing with acetate buffer, slices were rinsed several times with 0.1M sodium cacodylate buffer (pH 7.4) and post-fixed for 20 min in 1% osmium tetroxide supplemented with 1.5% ferricyanide in cacodylate buffer. Subsequently, the tissue was dehydrated using a gradient of 30%, 50%, 70%, 80%, 90% and 100% ethanol followed by acetone. After incubating for 30 minutes in a 1:1 mixture of acetone with epoxy, slices were incubated for 30 min in pure epoxy and left at 65°C overnight to harden. With an Ultracut UCT ultrathin 70 nm sections were made and collected on electron microscopy grids with a formvar film. Contrasting of the tissue was achieved by incubation with 0.5% uranyl acetate for 4 min, followed by extensive washing with milliQ and drying to the air, and subsequent incubation with lead citrate for 2 minutes. Ultrathin sections were examined with a FEI Tecnai G12 electron microscope (FEI, Europe NanoPort, Eindhoven, the Netherlands) and images obtained with a Veleta camera, acquired as 16-bits TIF files. Images were saved in tiff format and analyzed using Fiji (ImageJ). We examined > 100 sections from 3 mice/group.

#### Anti-Mouse MOG IgG ELISA

Serum was collected from animals at various timepoints following transfer to cuprizone diet and were kept frozen at –20°C until required. Samples were assayed using Anaspec SensoLyte ® Anti-Mouse MOG (1-125) IgG Quantitative ELISA kits (AS-55156). Briefly, 96-well plates precoated with recombinant mouse MOG protein (1-125) were incubated with 50 mL of the appropriate samples or standard with gentle shaking at RT for 1 hour. Each sample was diluted in sample buffer at 1:100 and subject to 1:4 serial dilutions up to 1:6400. Each sample was plated in duplicate on the precoated/preblocked plate. Following sample incubation, samples were washed 5 times with wash buffer and incubated with anti-mouse IgG-HRP (1:2000 dilution) with gentle shaking at RT for 1 hour. Following incubation with secondary antibody, the plate was washed 5 times and 100 mL of TMB was added to detect level of anti-MOG IgG via optical density at 450 nm using a spectrophotometer. Serum from hMOG-immunized EAE mice at the chronic phase of disease were used as a positive control for this assay and was assayed at a starting dilution of 1:100 subjected to 1:4 serial dilutions up to 1:1638400.

#### Statistical analysis

All tests were performed using GraphPad Prism software (versions 5 to 8, GraphPad Software Inc, San Diego, CA, USA). Sample sizes for the animal experiments and electrophysiological recordings were determined by performing power tests with a type II error set to 0.8. The type of variability of distributions were assessed by Shapiro-Wilk normality test. The non-normally distributed data was analysed with non-parametric Mann-Whitney test if two groups were compared or with the non-parametric Kruskal-Wallis test followed by Dunn’s correction for multiple comparisons if >2 groups were compared. Correlation analyses of non-normally distributed data was performed by Spearman correlation coefficient. If data were normally distributed data groups were analysed by either ordinary two-way or repeated measures (RM) parametric analysis of variance (ANOVA) followed by post-hoc analyses with Sidak’s or Dunnett’s correction for multiple comparisons. For all tests, the null hypothesis was rejected with *P* < 0.05 at a 95% confidence interval.

## Data availability

All raw data supporting the findings of this study are available from the corresponding authors upon reasonable request.

## Results

### CA2 enrichment of C1q deposits in the atrophic demyelinated MS hippocampus

To test for demyelination-dependent or -independent changes in C1q, the immunohistochemical analyses conducted in this study included both myelinated and demyelinated MS hippocampi. To this end, using a collection of post-mortem hippocampal tissue from 55 MS cases and 5 non-neurological controls (NNC), we first performed immunostaining for the PLP marker of myelin and identified 31 cases with myelinated, lesion-free, MS hippocampus (MS myelinated, MS-M) and 24 MS cases with partly or completely demyelinated hippocampus (MS demyelinated, MS-DM) (**Fig. 1a**). Consistent with previous work [28], the hippocampi from NNCs showed no sign of demyelination. In addition, and in line with previous reports [13, 34], the MS samples showed only a slightly increased HLA-DP-DQ-DR staining, suggesting enhanced microglial reactivity generally restricted to hippocampal areas in the context of preserved PLP myelin staining was preserved (data not shown).

**Figure 1.**
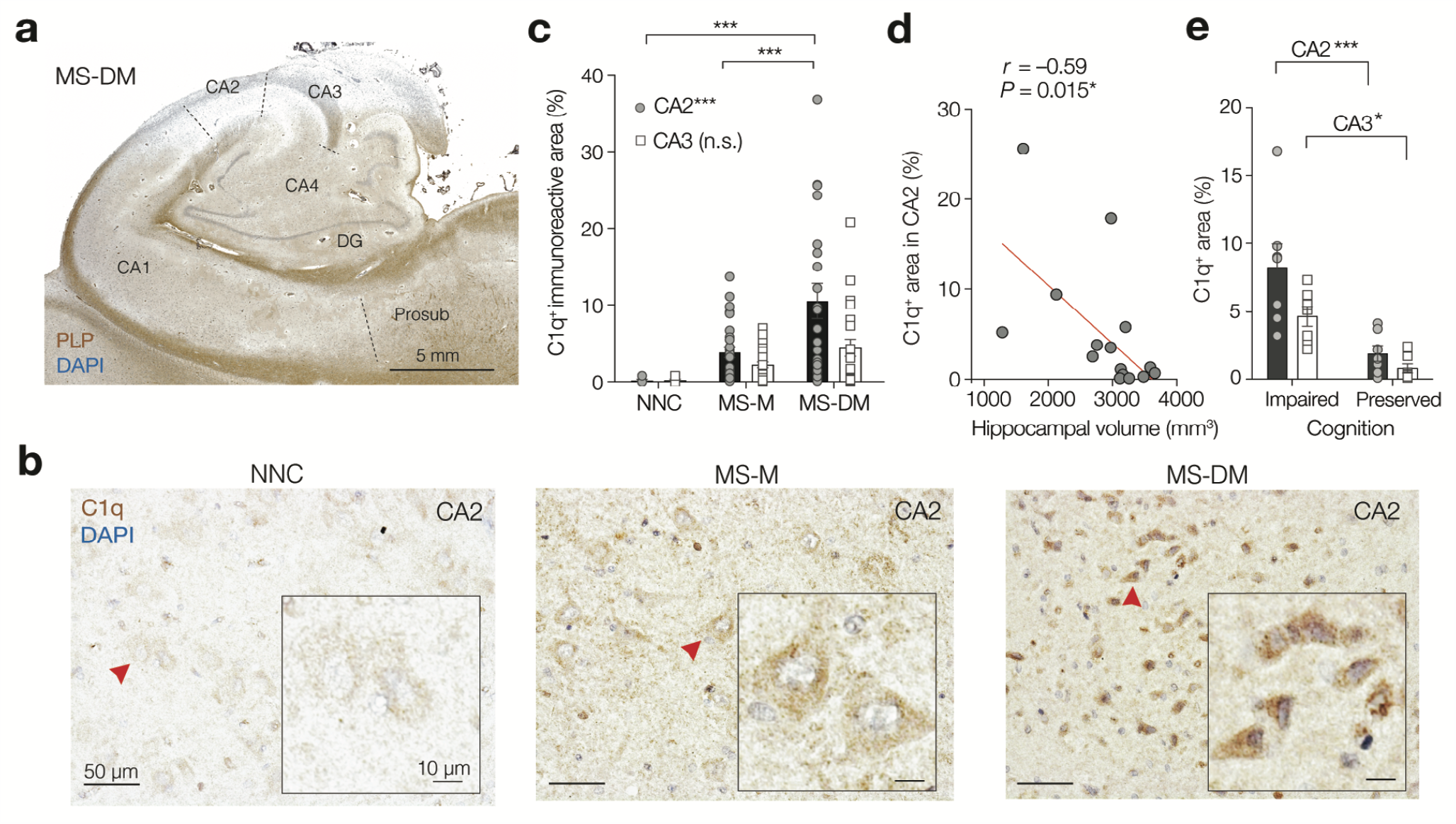
Multiple sclerosis hippocampus shows demyelination-dependent upregulation of C1q in the CA2. **a**. Coronal view of the hippocampus of a person with MS with hippocampal demyelination (MS-DM) in CA2 and CA3 assessed by proteolipid protein (PLP, brown) and DAPI (blue). **b**. Higher magnification of the CA2 subfield of hippocampi from a non-neurological control case (NNC), an MS case with myelinated hippocampus (MS-M) and a MS-DM case. Red arrows indicate location of the the insets showing individual CA2 neurons at higher magnification. Note the increased C1q immunoreactivity in perisomatic domains of MS-M and MS-DM compared to NNC. Decentrated neuronal nuclei are visible in the inset of MS-DM. **c**. Quantification of C1q immunoreactivity in post-mortem hippocampal CA2 and CA3 subfields. MS significantly increased C1q expression (two-way ANOVA *F*_2, 109_ = 11.36, *P* < 0.0001, NNC *n* = 5, MS-M *n* = 30, MS-DM *n* = 21), with a trend for subfield dependence (*F*_1, 109_ = 3.31, *P* = 0.0636). MS did not significantly upregulate C1q in the CA3 subfield (Sidak’s multiple comparison tests MS-M and MS-DM versus NNC, *t* = 0.74 and 1.53, respectively, *df* = 109 and n.s. for both). MS with hippocampal demyelination upregulated C1q in the CA2 area (For both MS–DM versus NNC and MS–DM versus MS-M; Sidak’s multiple comparison’s test, *t* = 3.72 and 4.21, respectively, \*\*\**P* < 0.0001). Bars represent the mean ± SEM; Grey circles and open squares represent individual hippocampi for CA2 and CA3 areas, respectively. **d**. Spearman’s correlation coefficient showing a significant negative correlation between C1q immunoreactivity in CA2 and hippocampal volume as determined by post-mortem MRI in a subcohort of MS cases (two-tailed exact *P* = 0.0145, *n* = 17 hippocampi). **e**. The intensity of C1q deposits was higher in MS donors with impaired cognitive/memory function compared to donors with preserved cognitive/memory function (two-way ANOVA cognition effect *F*_1, 24_ = 26.44, \*\*\**P* < 0.0001) and different between regions (two-way ANOVA subfield effect *F*_1, 24_ = 5.68, *P* = 0.0254, *n* = 7 biological replicates for all groups). C1q is higher in cognitively impaired MS patients in both CA2 and CA3 (Sidak’s multiple comparison test CA2, *t* = 4.55, *df* = 24, \*\*\**P* < 0.0001 and CA3, *t* = 2.73, *df* = 24, \**P* < 0.01, respectively). Bars represent the mean ± SEM; Circles and squares represent individual hippocampi for CA2 and CA3 areas, respectively.

Our previous work indicated that the amount of C1q immunoreactivity in the MS hippocampus was high in CA2/3 compared to other hippocampal subfields, including CA1 and subiculum [28]. Since it is becoming increasingly clear that the CA2 hippocampal subfield has a cytoarchitecture, connectivity, gene expression and neurochemistry functionally distinct from CA1 and CA3 [32, 35-37], we aimed to examine whether demyelination may have subfield-specific alterations. The C1q levels were determined by immunohistochemistry and the boundaries between CA3 and CA2 were based on cytoarchitectural criteria such as the higher cell packing density and larger CA2 pyramidal neurons somata [38]. Furthermore, because C1q expression in the CNS increases with normal ageing [27], we age-matched the donors to control for age-dependent changes in our samples. We confirmed the previously observed punctate staining pattern of C1q on and around hippocampal neurons, which was more obvious in MS cases compared to controls (**Fig. 1b**). In particular, C1q-coated CA2 neurons of MS-DM cases had a dystrophic appearance with decentrated nuclei, suggestive of neuronal injury (**inset, Fig. 1b**). Comparative analysis of the intensities showed that in the CA2 region of MS-DM hippocampi the amount of C1q deposition was on average ∼10-fold increased compared to NNCs and ∼3-fold compared to MS-M (*P* < 0.0001 for both, **Fig. 1c**) but not within CA3 (**Fig. 1c**). Because C1q deposits were enriched in CA2 compared to CA3 in the demyelinated MS hippocampus (Sidak’s multiple comparison test *P* < 0.0013) we next asked whether the density of C1q deposition in CA2 could be linked to hippocampal atrophy. Correlation analyses between the density of C1q staining and volumetric changes of the hippocampus as measured by post-mortem MRI revealed a significant correlation between the extent of C1q deposition and hippocampal volume (*P* = 0.015, **Fig. 1d**), demonstrating an association between C1q in CA2 and hippocampal atrophy in progressive MS donors. Furthermore, since the hippocampus is of critical importance for spatial and emotional memory, we next asked whether there is a link between C1q deposition in cognitive functions. Comparison of the immunofluorescence density of C1q in MS cases with or without documented impairment of cognitive function (based on available clinical records) showed that MS cases with impaired cognitive function had a significantly and 4.3-fold higher amount of C1q deposits in CA2 than those patients without evidence of cognitive problems (Sidak’s multiple comparison test *P* < 0.001, **Fig. 1e**). While the difference was also detectable in CA3 (Sidak’s test *P* < 0.05, **Fig. 1e**) the C1q expression was substantially higher within the CA2 region (two-way ANOVA, subfield *P* < 0.0254). Together with the significantly higher expression in the larger data set (**Fig. 1c**) these results indicate that the CA2 region shows an increased sensitivity to complement activation in MS.

### Loss of GABAergic and gain of glutamatergic synapse markers in the CA2 subfield of the MS hippocampus

While a common finding from our previous studies [28] and others [39, 40], is that synapses are lost in the MS hippocampus, which synapses are selectively changed within the CA2 hippocampal subfield is not well understood. We performed immunofluorescence staining for the presynaptic vesicular glutamate transporters 1 (vGLUT1) and a postsynaptic element of excitatory synapses the postsynaptic domain 95 (PSD95), as well as a presynaptic marker for gamma-aminobutyric acid (GABA)ergic synapses, the vesicular GABA transporter (vGAT), and a postsynaptic elements of inhibitory synapses (gephyrin). Quantification analysis of presynaptic elements in CA2 showed that compared to NNCs the amount of vGLUT1^+^ synapses was increased in MS tissue by 2.4-fold (One-way ANOVA *P* < 0.0001) while the amount of vGAT^+^ synapses in MS hippocampi was decreased 2-fold (One-way ANOVA *P* = 0.0267) (**Fig. 2a-d**). Similar changes were observed in the CA2 of MS-M cases (vGLUT^+^ puncta, 2.5-fold increase in MS-M vs NNCs, *P* < 0.01; vGAT^+^ puncta, 1.8-fold decrease in MS-M vs NNCs, *P* < 0.01, **Fig. 2a-d**). Furthermore, quantification analysis of postsynaptic elements in CA2 showed that compared to NNCs, the amount of PSD95^+^ synapses was increased 2-fold (*P* < 0.01) while, in striking contrast, the amount of gephyrin^+^ puncta decreased 7-fold in MS-DM hippocampi (*P* < 0.01, **Fig. 2**). Similar changes in gephyrin^+^ postsynaptic elements were observed in the CA2 of MS-M cases (6.6-fold decrease in MS-M vs NNCs, *P* < 0.01), whereas no changes were observed in PSD95^+^ postsynaptic elements of the CA2 of MS-M cases (**Fig. 2**). These findings indicate a gain of excitatory postsynaptic elements and a concomitant loss of inhibitory postsynaptic elements in the CA2 subfield of the MS hippocampus. Furthermore, they suggest that changes in inhibitory but not excitatory CA2 postsynaptic elements may precede demyelination in MS.

**Figure 2.**
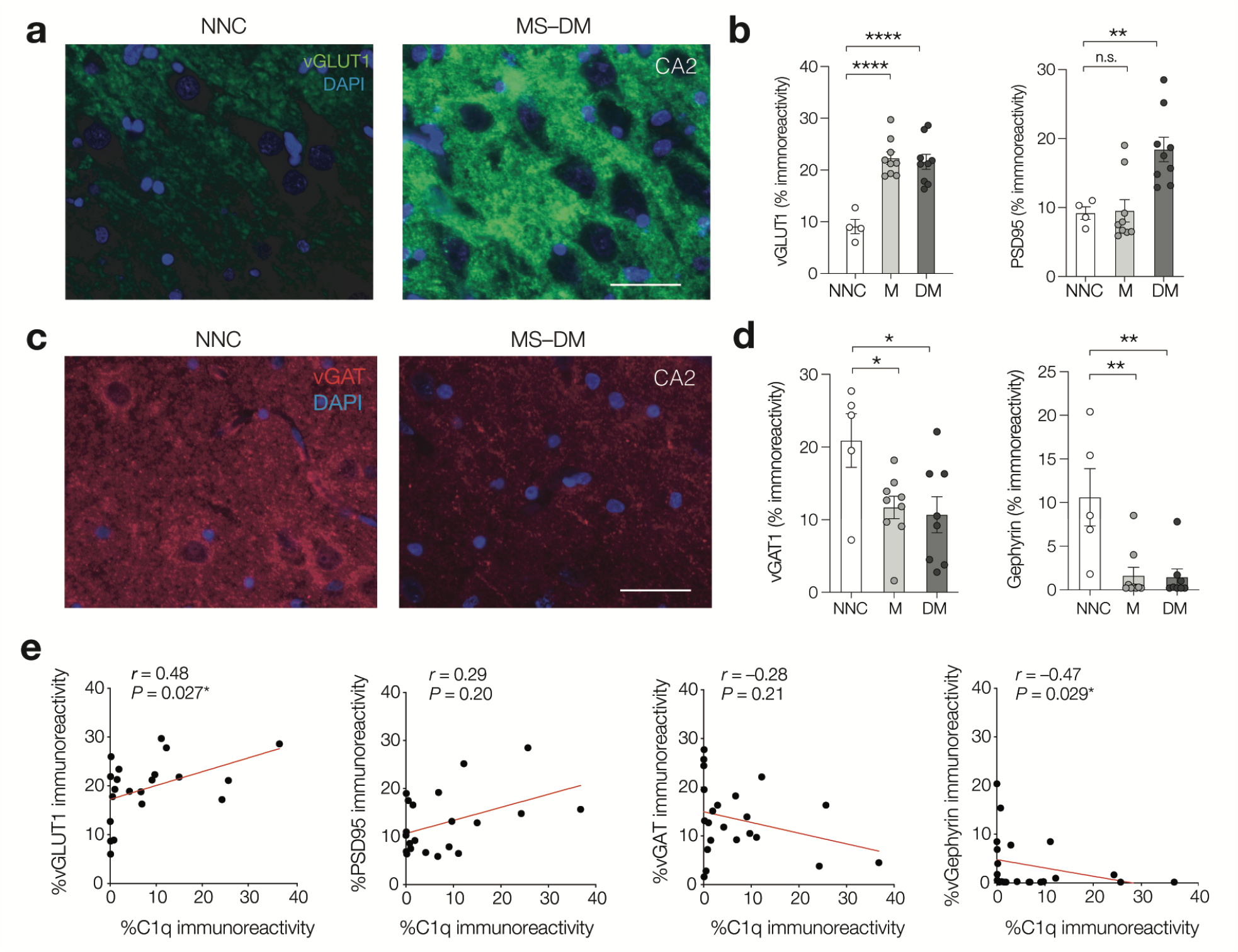
Diverse synaptic changes in the MS CA2 region associate with C1q expression. **a**. Immunofluorescence staining for vGLUT1 (in green) and DAPI (in blue) identifying glutamatergic synapses in CA2 stratum pyramidale of a non-neurological control (NNC) case and an MS case with demyelinated hippocampus (MS-DM). Scale bar, 20 µm. **b**. Population analysis of vGLUT1 and PSD95 in CA2 of non-neurological control (NNC) cases and MS cases with myelinated (MS-M) or demyelinated (MS-DM) hippocampi. MS increased vGLUT1 density (One-way ANOVA *F*_2, 19_ = 18.55, *P* < 0.0001, Dunnett’s multiple comparison test MS-M versus NCC, *q* = 5.75, *df* = 19, \*\*\*\**P* < 0.0001; MS-DM versus NCC, *q* = 5.44, *df* = 19, \*\*\*\**P* < 0.00001) and PSD95 (One-way ANOVA *F*_2, 19_ = 9.63, *P* = 0.0013, Dunnett’s multiple comparison tests, MS-M versus NCC, *q* = 0.11, *df* = 19, *P* = 0.998; MS-DM versus NCC, *q* = 3.24, *df* = 19, \*\**P* = 0.0075). **c**. Immunofluorescence staining for vGAT (in red) and DAPI (in blue) identifying GABAergic synapses in CA2 stratum pyramidale of a non-neurological control (NNC) case and an MS case with demyelinated hippocampus (MS-DM). Scale bar, 20 µm. **d**. Population analysis of vGAT and gephyrin in CA2 in NNC, MS-M and MS-DM cases. MS is associated with a loss of inhibitory presynaptic component vGAT (One-way ANOVA *F*_2, 19_ = 4.41, *P* = 0.0267, followed by Dunnett’s tests; MS-M versus NNC, *q* = 2.56, *df* = 19, \**P* = 0.034; MS-DM versus NNC, *q* = 2.78, *df* = 19, \**P* = 0.021) as well as a loss of inhibitory postsynaptic density protein gephyrin (One-way ANOVA *F*_2, 19_ = 9.18, *P* = 0.0016) both for DM and M cases (Dunnett’s multiple comparison tests *q* = 3.87, *df* = 19, \*\**P* = 0.0019 and *q* = 3.86, *df* = 19, \*\**P* = 0.0019, respectively). **e**.Correlation analyses between the amount of C1q and the amount of vGLUT1^+^ or PSD95^+^ or VGAT^+^ or gephyrin^+^ synapses in CA2 of MS cases (*n* = 17). Spearman’s correlation coefficient (*r*) shows a significant positive correlation between the amount of C1q and the amount of vGLUT-1^+^ synapses (*n* = 21) whereas it shows a significant negative correlation between the amount of C1q and the amount of gephyrin^+^ synapses (*n* = 22). Two-tailed *P* values indicated in the figure panels.

### C1q deposits correlate with synaptic changes in the CA2 subfield of the MS hippocampus

To determine whether there is a link between the density of C1q deposition and synaptic changes in the MS hippocampus, we next performed correlation analyses between the density of C1q staining and either the density of vGLUT1^+^ or vGAT^+^ or PSD95^+^ or gephyrin^+^ synapses determined in the same MS hippocampi. Combining the control, myelinated and demyelinated MS hippocampi, we found a significant correlation between the extent of C1q deposition and the density of vGLU1^+^ synapses (Spearman correlation coefficient, *r* = 0.48, *P* = 0.027, *n* = 21), as well as between the extent of C1q deposition and gephyrin^+^ synapses (Spearman correlation coefficient, *r* = –0.47, *P* < 0.029, *n* = 22), indicating an association between C1q, gain of excitatory synapse markers and loss of inhibitory synapses in the CA2 subfield of the MS hippocampus (**Fig. 2e**). Although the extent of C1q deposition increased with decreasing vGAT^+^ synapses (*r* = –0.28) and with increasing PSD95^+^ synapses (*r* = 0.28), these associations did not significantly correlate (*P* = 0.2 for both, **Fig. 2e**). Taken together, these data demonstrate an association between C1q and synaptic reorganization in the CA2 hippocampal subfield of progressive MS donors.

### Enrichment of C1q in the dorsal CA2 subfield in the demyelinated hippocampus of cuprizone-fed mice

To understand the role of myelin loss and determine the functional consequences of C1q-mediated synapse loss in the CA2 subfield we next investigated the hippocampus in the cuprizone mouse model [41, 42]. Sagittal slices were cut along the dorsal-to-ventral axis of the hippocampus of adult (4-months old) male mice and stained for myelin basic protein (MBP) and compared with age-matched mice treated with cuprizone (0.2% for 9 weeks, **Fig. 3a**). In the control hippocampus, MBP was densely distributed in the white-matter tracts (fimbria and alveus) and the perforant path. In addition, MBP was also observed throughout the intrahippocampal grey matter regions including CA3 and CA2 (**Fig. 3a**). Consistent with previous studies with cuprizone [43-46], myelin was strongly reduced in the white matter regions including the alveus and fimbria and near completely lost in the intrahippocampal grey matter areas (**Fig. 3a**). This pattern of intrahippocampal myelin loss was highly reproducible across mice and observed along the entire dorsal-to-ventral hippocampal axis (**Supplementary Fig. 1**).

**Figure 3.**
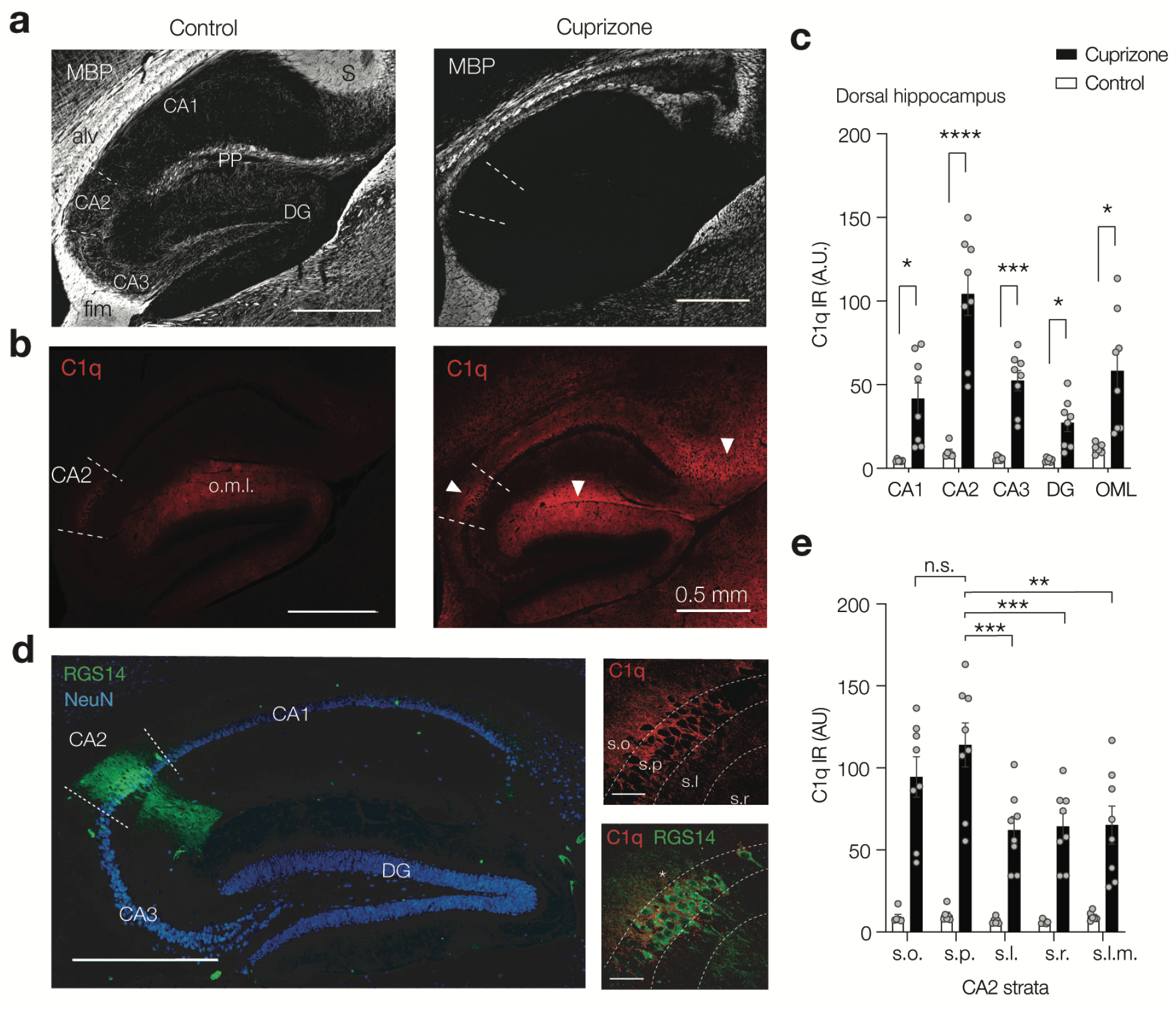
Cuprizone-induced demyelination causes subregion dependent C1q increase. **a**. *Left*, Example fluorescence image of a sagittal section of the control dorsal hippocampus for myelin basic protein (MBP-Ab, white) and, *right*, following 9 weeks 0.2% cuprizone treatment. In control, high intensity signals are present in the white-matter tracts including the alveus (alv) and fimbria (fim) and the myelinated perforant path (PP) fibers. Lower intensity signals are present in the CA3 and CA2 stratum pyramidal and oriens layer but also in the dentate granule (DG) moleculer layers. Note the loss of intra-hippocampal MBP expression and intrahippocampal differences in MBP distribution with low levels in CA1 but relatively stronger signals in CA2 and CA3, as well as the and molecular layers of the DG. Scale bar, 0.5 mm. **b**. *Left*, control expression of complement factor 1q (C1q-Ab, red). Low levels are visible in CA2 as well as the DG outer molecular layer (o.m.l.). *Right*, demyelination increases C1q (red) widely across hippocampal subfields and with high intensities in CA2, molecular layers and subiculum (white arrows). **c**. Population analysis of C1q intensity in the dorsal hippocampus shows C1q immunoreactivity increases in a subregion-specific manner following cuprizone treatment (two-way RM ANOVA, Treatment *F*_1, 12_ = 23.71, *P* = 0.0004; Subregion *F*_2180, 26.16_ = 30.51; *P* < 0.0001 and Subregion × Treatment *F*_4, 48_ = 23.02, *P* < 0.0001, *n* = 6 sections from 3 animals/group). After cuprizone treatment the CA2 region shows the highest intensities in comparison to CA1, CA3, the outer molecular layer and the DG (Sidak’s multiple comparisons test, CA1 \**P* = 0.0261, CA2 \*\*\**P* = 0.0007, CA3 \*\*\**P* = 0.0007, DG \**P* = 0.0157, DG \**P* = 0.0357). Post-hoc test for regions revealed C1q within CA2 was higher in comparison to all other subregions (Sidak’s multiple comparisons test \*\*\**P* < 0.0001, data not shown). **d**. *Left*, overview image of the hippocampus with the CA2-specific marker, anti-regulator of G protein signaling 14 (anti-RGS14, green), staining somata and dendrites of CA2 pyramidal neurons and anti-NeuN (blue). Scale bar, 500 µm. *Right*, higher-magnification images reveal C1q clusters in the perisomatic of CA2 neurons (RGS14, green). One RGS14^−^ neuron with perisomatic C1q indicated with an asterisk. Stratum oriens (s.o.), stratum pyramidale (s.p.), stratum lucidum (s.l.), stratum radiatium (s.r.) and stratum lacunosum-moleculare (s.l.m.). Scale bars, 50 µm. **e**. Population analysis of C1q intensity across the distinct strata within CA2 reveals a strata-specific cuprizone-induced C1q increase (2-way ANOVA Treatment *F*_1, 60_ = 156.8, *P* < 0.0001, Strata *F*_4, 60_ = 3.69 *P* = 0.0094 and Treatment × strata *F*_4, 60_ = 2.95, *P* = 0.0281, *n* = 8 sections from 4 mice) with after cuprizone treatment s.p. showing higher C1q intensity compared to other strata (Sidak’s multiple comparison tests, versus s.l.m.\*\**P* = 0.0012, s.r. \*\*\**P* = 0.0009, ***s.luc *P* = 0.0005). However, C1q intensities in s.p. and s.o. were similar, *P* = 0.668). Error bars indicate mean ± SEM and grey dots individual section.

Immunofluorescence staining for C1q was performed in the same sections that were also stained for MBP. Consistent with previous reports [27, 47], in the control hippocampus low intensities of C1q immunoreactivity was already detected specifically within the CA2 subfield and the molecular layer of the dentate gyrus (DG) (CA2 and DG o.m.l., **Fig 3b**), Following cuprizone feeding, however, C1q immunoreactivity increased widely across the intrahippocampal grey matter and parahippocampal white-matter regions (on average 7.8-fold, *P* < 0.0001, **Fig 3b, c**). Quantitative immunofluorescence analysis across the different hippocampal subfields showed that average C1q intensities were significantly higher in CA2 compared to CA1, CA3 and the DG (**Fig. 3c**). Furthermore, the cuprizone-induced upregulation of C1q followed a gradient along the longitudinal axis with the highest expression level in the dorsal region, lower in the intermediate region and undetectably low in the ventral hippocampus (**Supplementary Fig. 1a-d**). To further investigate the prominent CA2 localization of C1q in the demyelinated hippocampus, we stained hippocampal slices from control and cuprizone-fed mice with RGS14, a specific molecular marker for CA2 pyramidal neurons [48]. Co-staining for RGS14 and C1q showed that C1q was predominantly clustered in the stratum pyramidale and oriens of RGS14^+^ neurons at significantly higher intensities when compared to the stratum lucidum, radiatum and lacunosum moleculare (**Fig. 3d, e)**. Interestingly, a few RGS14^−^ neurons in CA2, presumably interneurons, also showed perisomatic C1q (**Fig. 3d, e**).

Thus, in accordance with the demyelinated MS hippocampus (**Fig. 1**) C1q localization is highly circuit specific (CA2) and cuprizone-induced demyelination causes a strong enrichment around the CA2 pyramidal (RGS14^+^) neurons.

In the MS hippocampus (**Fig. 1** and [28]), models of neurodegeneration [22] and EAE models of demyelination[16] some C1q expression is already detectable before overt signs of pathology or myelin loss. We thus next asked whether the increase in C1q deposition at CA2 precedes or follows the loss of myelin in this region. We quantified the extent of C1q immunoreactivity in relation to MBP immunoreactivity in the CA2 subfield throughout the course of cuprizone feeding (up to 6 weeks). The C1q immunoreactivity increased about 2-fold from baseline levels at 2 weeks of cuprizone feeding and reached its maximum around 4 weeks. These changes in C1q were mirrored by a loss of MBP immunoreactivity in the CA2 subfield with levels rapidly decreasing by 2-fold at 2 weeks of cuprizone feeding and reaching maximum loss around 6 weeks (**Supplementary Fig. 2a, b**). Correlation analysis showed a significant negative correlation for the CA2 stratum pyramidale layer (*r* = –0.66, *P* < 0.0001, **Supplementary Fig. 2c**) supporting a link between the loss of myelin and the C1q increase in CA2 in this model.

Since a classical role of C1q is to tag antigen/(auto)antibody complexes for elimination[49], and anti-myelin antibodies are detected in serum of models of (auto)antibody-mediated demyelination[50], we next examined whether cuprizone-induced upregulation of C1q in CA2 was associated with serum titers of anti-myelin antibodies at the time when we detect myelin loss in the hippocampus. To test this, we measured anti-myelin oligodendrocyte glycoprotein (MOG) antibody levels in serum from these mice throughout the 6 weeks of cuprizone feeding as well as C57BL/6 mice immunized with human recombinant MOG (hMOG) protein as a technical positive control because hMOG-immunized C57BL/6 mice generate anti-MOG IgG antibodies that are required for the manifestation of clinical disease[50]. As expected, anti-MOG IgG titers were detected in hMOG-immunized C57BL/6 mice. In addition, anti-MOG IgG titers were very low or not detected in control mice (as expected) or cuprizone-fed mice throughout the 6 weeks of cuprizone feeding, including time points when myelin loss is evident (**Supplementary Fig. 2d**). These findings indicate that anti-MOG antibodies are not required for the observed demyelination and are unlikely to be the trigger of C1q upregulation in tissue. In summary we show that in the mouse hippocampus, cuprizone feeding triggers a circuit- and cell-specific increase in C1q immunoreactivity that is strongly linked with demyelination but is independent of anti-MOG antibodies, suggesting an antibody-independent role of C1q in the CA2 hippocampal subfield.

### Gain of excitatory synapses but loss of inhibitory synapses in the CA2 hippocampal subfield

The perisomatic clustering of C1q in CA2 stratum pyramidale caused by cuprizone-induced demyelination is strikingly consistent with the observations in the human MS hippocampus (**Fig. 1**). To determine the specific synaptic changes in the demyelinated CA2 we used double immunofluorescence staining for RGS14 (to identify CA2) in combination with synaptic markers that identify either glutamatergic synapse including the postsynaptic protein Homer1 and the presynaptic vGLUT1 and vGLUT2 (**Fig. 4a**) or vGAT (**Fig. 4b**). Quantification of puncta within CA2 revealed a significant increase in the density of excitatory (vGLUT1^+^, vGLUT2^+^ and Homer1^+^) synapses and a concomitant significant reduction in the density of inhibitory (vGAT^+^) synapses (**Figs. 4a-c**). Notably, unlike the vGLUT1 puncta, which were widely distributed in strata radiatum and oriens, the vGAT^+^ synapses were clustered in the strata oriens and pyramidale, where the highest immunoreactivity for C1q was present and vGAT^+^ synapses apparently contacted CA2 pyramidal neuron cell bodies (**Figs. 4b**). To examine whether C1q localizes at synapses in the CA2 region during cuprizone-induced demyelination we performed triple labelling for C1q, RGS14 and synapse markers. We found that compared to controls, cuprizone treatment significantly increased C1q^+^/vGLUT2^+^ and C1q^+^ /vGAT^+^ synaptic contacts, but not C1q^+^/vGLUT1^+^ synapses (**Fig. 4d-f**). Population analysis showed that there was a significant >3-fold increased probability that vGLUT2 and vGAT synapses were in contact with C1q and arranged similarly around the CA2 soma (**Fig. 4d-f**).

**Figure 4.**
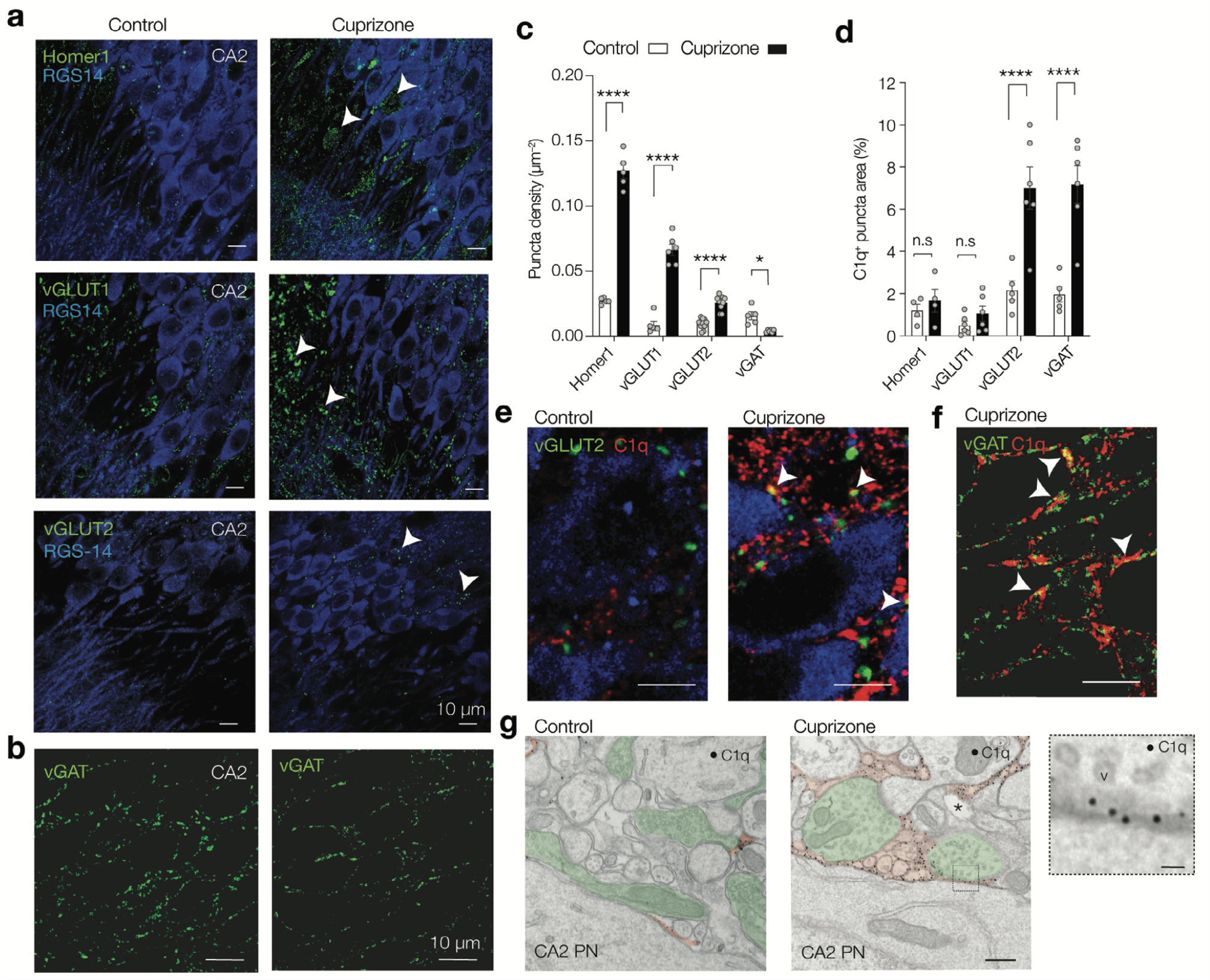
Bidirectional change in excitatory and inhibitory synapse markers in CA2. **a**. Example immunofluorescent staining for RGS14 (blue) and postsynaptic glutamate receptor Homer1 (top, green), the presynaptic excitatory vesicular glutamate transporter 1 (vGLUT1, middle, green) and vGLUT2 (bottom, green) in control (left) and cuprizone hippocampus (right). Note the increase vGLUT1 puncta in the strata lucidum and radiatum and localization of vGLUT2 in pyramidale and oriens layers. White arrows indicate example puncta included in counting. Scale bar, 10 µm. **b**. Example images of vesicular GABA transporter (vGAT, green) overlaid with DAPI (cyan) in the CA2 region of control (left) and cuprizone hippocampus (right). Note the loss in vGAT puncta. Scale bar, 10 µm. **c**. Population data for synaptic marker densities reveals a gain in excitatory-but loss of inhibitory synapse markers (Two-way ANOVA Treatment effect *F*_1, 45_ = 427.3, *P* < 0.0001, Treatment × Synapse marker interaction *F*_2, 45_ = 139.4, *P* < 0.0001, followed by Sidak’s multiple comparison test for Homer1 (*t* = 22.95, *df* = 45, \*\*\*\**P* < 0.0001), vGLUT1 (*t* = 14.56, *df* = 45, \**\*\*\*P* < 0.0001), vGLUT2 (*t* = 5.25, *df* = 45, \*\*\*\**P* < 0.0001) and vGAT (*t* = 2.80, *df* = 45, \**P* = 0.0297). Each group represents *n* = 5–10 sections from 6 control and 4 cuprizone-treated mice. **d**. Population data for C1q co-localization (% area overlap) in control (open bars) and cuprizone-treated mice (closed bars). Cuprizone-induced increase in C1q is differentially distributed across synapses (Two-way ANOVA treatment × synapse *F*_3, 34_ = 8.64, *P* = 0.0002) and significantly co-localizes to vGLUT2 and vGAT markers (Sidak’s multiple comparison test *t* = 5.57, *df* = 34, vGLUT2, \*\*\*\**P* < 0.0001, *n* = 5 control and 6 cuprizone, and vGAT *t* = 6.00, *df* = 34,\*\*\*\**P* < 0.0001, *n* = 5 control and 6 cuprizone) but not Homer1 nor vGLUT1 markers (Sidak’s multiple comparison tests, Homer1; *t* = 0.46, *df* = 34 *P* = 0.985, *n* = 4 both groups and vGLUT1; *t* = 0.66, *df* = 34, *P* = 0.945, *n* = 6 both groups). Data represented as mean + SEM with individual sections indicated with circles. **e**. Example triple immunostaining images for RGS14 (blue), C1q (red) and vGLUT2 (green) in control and cuprizone hippocampus. White arrows indicate co-localization of C1q and vGLUT2 (yellow, white arrows). Scale bar, 10 µm. **f**. Double immunostaining for vGAT (green) and C1q (red). Note the perisomatic localization of vGAT and co-localization with C1q (yellow color, white arrows). Scale bar, 10 µm. **g**. Transmission EM images in the perisomatic region of a CA2 pyramidal neuron (CA2 PN) of a control (left) and a cuprizone-treated mouse (right). The anti-C1q immunogold (∼10 nm black particles, false colored red) are predominantly in the extracellular space near synapses (false colored green), both near putative inhibitory or excitatory synapses (right image, note the asymmetric postsynaptic density and spine). Scale bar, 400 nm. Right inset, higher magnification of a putative inhibitory synapse at the CA2 soma with C1q-IR gold particles and synaptic vesicles (v). Scale bar, 50 nm.

Since activation of the classical complement pathway, initiated by the binding of C1q to its target, results in activation of the downstream complement component C3, and since C3 has been involved in elimination of synapses during development [18] and in the MS visual thalamus [16], we next investigated whether also C3 activation products, like C1q, are deposited at synapses in the cuprizone hippocampus. Immunofluorescence staining for the membrane bound product of C3 activation, C3d, showed a significant 1.8-fold increase in C3d deposits in CA2 of cuprizone mice compared to controls. Interestingly, C3d co-localized with GFAP^+^ astrocytes (**Fig. S3a, b**), which may reflect the neurotoxic A1 type of astrocytes previously described in MS tissue [51]. C3d also co-localized with some synapses, however the amount of C3d^+^ synapses did not vary between cuprizone and control mice (**Fig. S3c, d**). Finally, immunogold electron microscopy (EM) for C1q protein confirmed that C1q is present at a low density in control CA2 subfield [27] but strongly increased throughout the extracellular spaces and matrix, often in close proximity to presynaptic terminals (**Fig. 4g**). Together, these data suggest that cuprizone feeding may cause a C1q-mediated reorganization of synapses in the CA2 pyramidal and oriens layers.

### C1q-tagged synapses localize within microglia/macrophages in the CA2 hippocampal subfield during cuprizone-induced demyelination

Complement-tagged synapses are eliminated via phagocytoses by microglia during development, adulthood, normal ageing, neurodegeneration and demyelination of the visual thalamus [16, 18, 22, 26, 27]. To test whether microglia/macrophages engulf C1q-tagged synapses in CA2, we first quantified changes in the number of cells positive for ionized calcium-binding adaptor molecule 1 (Iba-1), a marker for microglia/macrophages. Quantification of Iba-1^+^ cells in CA2 showed a significant 2-fold increase in number as well as a significant 3-fold increase in the area covered by the Iba-1^+^ cells in the cuprizone-treated mice compared to controls (**Fig. 5a, b)**. Double immunofluorescence labeling of Iba-1 with either Homer 1, vGLUT1, vGLUT2 or vGAT showed a basal level of co-localization of Iba-1 with each of the synaptic markers in the control hippocampus, likely reflecting ongoing surveillance of microglia. However, cuprizone feeding induced a specific increase in Iba-1^+^/vGLUT2^+^ and Iba-1^+^/vGAT^+^ synapses (**Fig. 5c-e)**. 3D rendering also showed microglial/macrophage processes surrounding synapses (see example of vGAT^+^ synapse in **Fig. 5d**), pointing to the engulfment of synaptic elements by myeloid cells in the CA2 area. In line, immunogold-EM for C1q protein in cuprizone-fed mice showed thick microglial processes engulfing electrodense element in close proximity to synapses (**Fig. 5f)** or touching C1q labeled synapses in the CA2 hippocampal subfield (**Fig. 5g)**, further supporting the close association and engulfment of synaptic elements by myeloid cells in the CA2 area.

**Figure 5.**
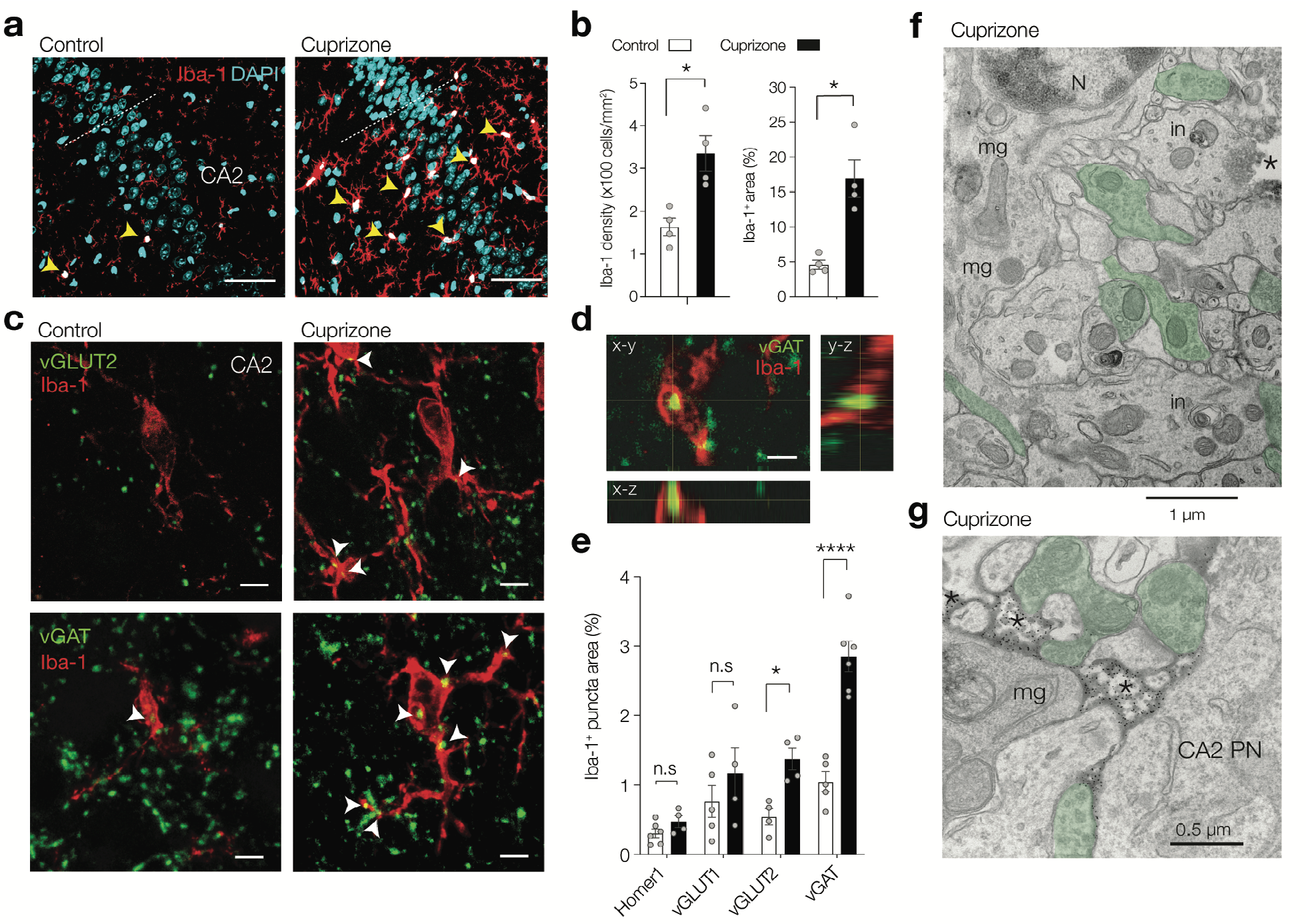
Activated microglia preferentially target vGLUT-2 and vGAT synapses in the CA2 region. **a**. Cuprizone increases microgliosis (anti-Iba-1, red) in the hippocampal CA2 subfield, identified with nuclear DAPI stain (blue). Note the increased size of microglia/macrophages (yellow arrows). Scale bar, 50 µm. **b**. Population analysis of the percentage of DAPI and Iba-1 positive cells (DAPI^+^ Iba-1^+^, white) within area CA2 shows cuprizone doubles the number of Iba-1^+^ cells (two-tailed Mann-Whitney test *P* = 0.0286) and increases the surface area by ∼3-fold (two-tailed Mann-Whitney test *P* = 0.0286, *n* = 4 sections from 4 animals). **c**. Double immunostaining for Iba-1 and vGLUT2. Note the increased overlap between Iba-1 and vGLUT2 and vGAT (white arrows) in the face of a loss of vGAT puncta. Scale bar, 5 µm. **d**. Higher magnification and orthogonal view of a putatively engulfed vGAT^+^ terminal (green) by Iba-1 (red). Same synapse as indicated by white arrow in c. Scale bar, 2 µm. **e**. Population data for the overlap of area between Iba-1 and synaptic markers Homer1, vGLUT1, vGLUT2 and vGAT shows a significant cuprizone treatment-induced preference of microglia contact with vGLUT2 and vGAT (Two-way ANOVA Treatment × Synapse marker *F*_3, 30_ = 7.81 *P* = 0.0005, Treatment *F*_1, 30_ = 34.17, *P* < 0.0001, followed by Sidak’s multiple comparison tests, Homer1 *t* = 0.633, *df* = 30, *P* = 0.952; vGLUT1 *t* = 1.42, *df* = 30, *P* < 0.512; vGLUT2 *t* = 2.81, *df* = 30, \**P* = 0.0337; vGAT *t* = 7.18, *df* = 30, \*\*\*\**P* < 0.0001, for all *n* = 4–6 sections from *n* = 4 animals/group). **f**. EM of an activated microglia (*mg*) in the CA2 region. The microglia nucleus (*N*) is identified by clumped chromatin. Note the cell body extends a thick large process towards presynaptic terminals (false green colored). Microglia processes contained an inclusion with phagocytosed debris (*In*), lysosomes, golgi apparatus, mitochondria and ER. **g**. Higher magnification of C1q-immunogold EM showing a microglia process with darker cytoplasm (*mg*) in the vicinity of a CA2 PN. C1q containing regions (black asterisks) are near presynaptic terminals (false green colored).

### Reduced feedforward inhibition of CA2 pyramidal neurons

What are the functional consequences of microglia/macrophages phagocytosed and C1q-tagged synapses for information processing in the CA2 circuit? The CA2 PNs receive a strong excitatory drive from layer 2 medial and lateral entorhinal cortex pyramidal neurons at their distal dendrites in the lacunosum moleculare but weak excitation from the DG mossy-fibers at the proximal dendrites [35, 52-54]. In the stratum pyramidale and oriens layers, where C1q immunoreactivity was markedly increased (**Figs. 4** and **5**), vGLUT2 reflects glutamatergic innervation from the medial septum diagonal band complex and the hypothalamic supramammillary nucleus [55, 56], whereas the vGAT are predominantly from fast-spiking parvalbumin (PV)^+^ interneurons producing feedforward inhibition from the CA3 Schaffer collateral (SC) axons [53, 54, 57, 58] (**Fig. 6a**). The CA2 PV^+^ interneurons are furthermore subjected to neuromodulation from the hypothalamic paraventricular nucleus (PVN) and supraoptic nucleus (SON), playing a critical role in encoding social learning by affecting plasticity of PV interneurons [57-59]. To test whether loss of vGAT synapses causes functional changes in the local inhibitory circuit of CA2 we electrically stimulated SC axons while recording in whole-cell patch-clamp configuration from CA2 PN somata in transverse slices from the dorsal hippocampal region from control and cuprizone-fed mice (6 weeks for 0.2% cuprizone, **Fig. 6a**). All recorded neurons were simultaneously filled with biocytin and post-hoc stained with RGS14 or PCP4. About 80% of the recorded neurons (*n* = 29/36) were unequivocally CA2 pyramidal neurons and included for further analysis for their properties.

**Figure 6.**
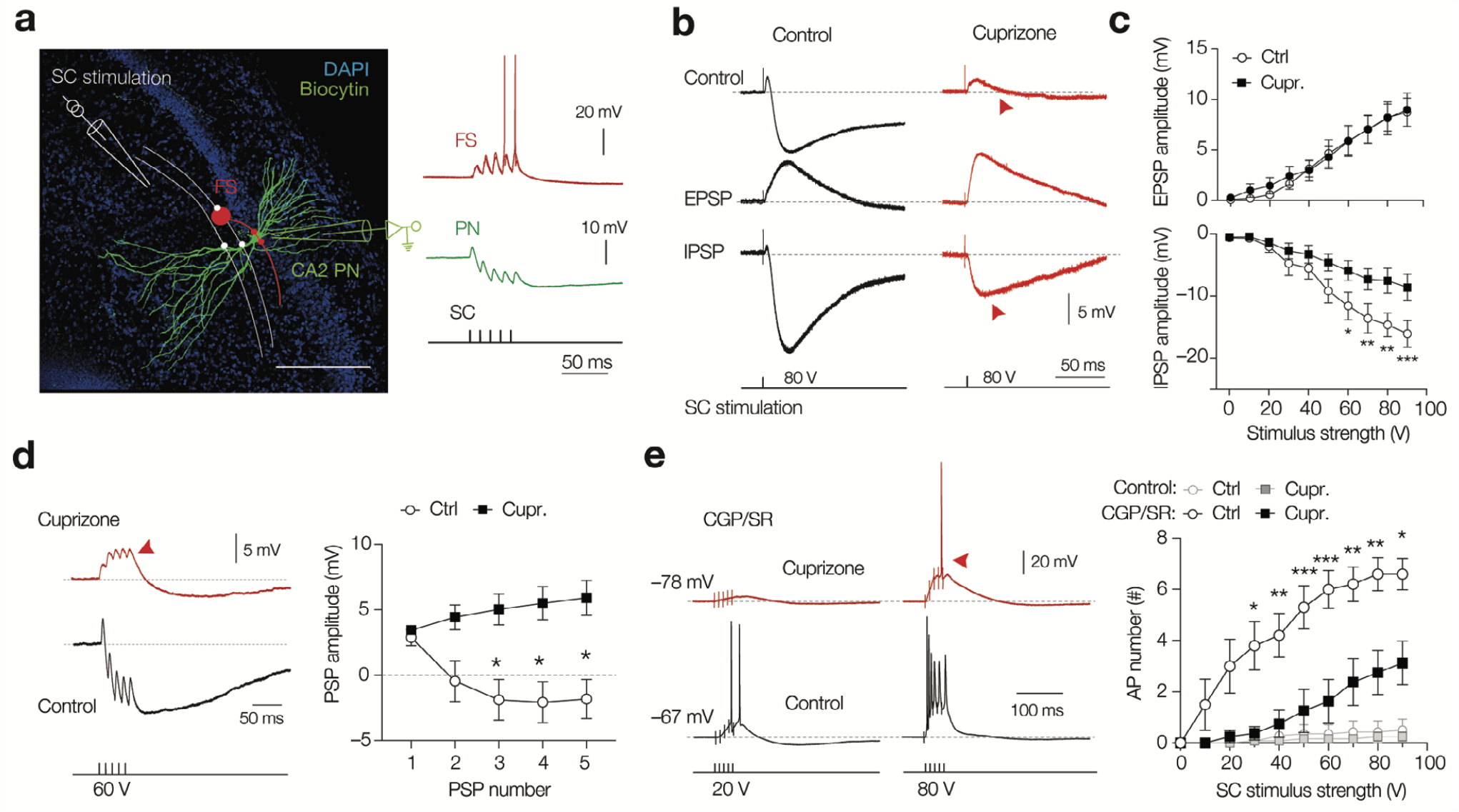
Cuprizone causes a loss of CA3 to CA2 feedforward inhibition and synaptically-evoked spike generation. **a**. Left, Immunofluorescence image of a biocytin-recovered CA2 pyramidal neuron (CA2 PN, green) and DAPI (blue) overlaid with schematic of a stimulation pipette for Schaffer collateral commissural axon fibers (SCs, white). Right, whole-cell recordings of a fast spiking (FS) interneuron showing typically strong temporal summation and SC-evoked spike output (red). In contrast, in CA2 PNs excitation is shunted by strong feedforward inhibition (green). SCs were activated with 5 × 60 µV pulses (100 Hz, 0.3 ms duration). **b**. Top, SC-evoked potentials in the CA2 PN from control (grey traces) and cuprizone (red traces). Middle, same recordings after bath application of the GABAergic antagonists CGP and SR, isolating excitatory postsynaptic potentials (EPSP, light gray). Bottom, digitally subtracted traces (control–EPSP) revealing the underlying IPSP. Note the reduced amplitude of the IPSP in CA2 PNs in slices from cuprizone-treated mice (red arrow). **c**. Population data for isolated EPSPs and IPSP as a function of stimulus strength (0–90 µV). The EPSPs amplitudes were unaffected by cuprizone treatment (mixed-effects model RM ANOVA, Treatment *F*_1, 11_ = 0.062 *P* = 0.809, *n* = 5 control and 8 cuprizone neurons from 6 mice/group). In contrast, IPSP peak amplitudes were significantly reduced (mixed-effects model RM ANOVA Treatment *F*_1, 13_ = 8.71, *P* = 0.0112, Treatment × stimulus interaction *F*_9, 96_ = 4.95, *P* < 0.0001, *n* = 8 neurons from 6 mice/group) and increased at stimulus intensities > 60 V (Sidak’s multiple comparison test 50 V *P* = 0.071, 60 V \**P* = 0.0146, 70 V \*\**P* < 0.0053, 80 V \*\**P* = 0.0017 and 90 V \*\*\**P* = 0.0010) **d**. Left, example traces of SC-evoked postsynaptic potentials, reflecting small excitation followed by feedforward inhibition (control) at 100 Hz train of subthreshold 60 V stimuli. Note the strong accumulation of inhibitory potentials in control CA2 neurons but not in cuprizone neurons (red arrow). Right, population data of the average peak amplitude responses (positive deflection relative to resting potential) as a function of stimulus pulse number (2-way RM ANOVA stimulus *F*_1244, 25_ = 2.53, *P* = 0.12, Treatment *F*_20, 80_ = 16.26, *P* = 0.0076, Treatment × stimulus interaction *F*_4, 80_ = 8.477, *P* < 0.0001) with significantly increased amplitudes for the 3^rd^ to 5^th^ stimulus (For all, Sidak’s multiple comparisons test *P* < 0.05, *n* = 13 control and 9 cuprizone neurons from 6 mice/group). **e**. Left, example traces for SC evoked excitation at 100 Hz in the presence of CGP 35348 and SR 95531. Note the low spike probability in the recordings from cuprizone treatment (top, red traces). Population data summarizing AP number per train across the range of stimuli in physiological extracellular solution (control, gray lines and symbols) or with blocked inhibition (CGP/SR, black open and closed symbols). Cuprizone suppressed SC-mediated CA2 PN spike output in stimulus strength dependence (two-way ANOVA Treatment *F*_1, 15_ = 14.79, *P* = 0.0016 Treatment × stimulus *F*_9, 135_ = 4.183, *P* < 0.0001).

In PNs in control slices, SC activation produced a brief depolarization followed by a strong inhibitory potential (**Fig. 6a**,**b**, [52]). A single brief SC stimulus was examined with varying levels of output voltage (0 to 90 V). The depolarizing peak of the postsynaptic potential (PSP) did not differ between the two groups across the range of stimuli (mixed-effect ANOVA Treatment *F*_9,171_ = 0.688, *P* = 0.548, at 90 V; control PSP amplitude, 3.20 ± 0.59 mV, *n* = 12, vs. cuprizone PSP 4.04 ± 0.64, *n* = 9, **Fig. 6b**, data not shown). To distinguish between the monosynaptic glutamate receptor and disynaptic GABAergic receptor activation in the Schaffer collateral pathway we applied CGP 35348 (antagonist of GABA_B_ receptors, 20 µM) and gabazine (SR 95531, a selective GABA_A_ antagonist, 3 µM). Analysis of the remaining EPSP revealed that SC activation in cuprizone-treated mice produced similar peak amplitudes across the entire range of stimuli (mixed-effect RM ANOVA, Treatment *P* = 0.809, **Figs. 6b, c**). In contrast, the IPSP peak amplitudes (subtracting the control PSP with the EPSP), were significantly reduced in a stimulus-dependent manner (**Figs. 6b, c**). Calculating the relative contribution within experiments (EPSP/IPSP amplitude ratios) confirmed a significant loss of inhibition of CA2 PNs from cuprizone-treated mice (control ratio, 0.64 ± 0.06 versus cuprizone ratio 1.11 ± 0.15, two-tailed Mann-Whitney test U = 7, *P* = 0.0064, *n* = 8 neurons from 6 mice/group). About ∼80% of the GABAergic cells in the CA2 stratum pyramidal layer are parvalbumin-positive (PV^+^) interneurons[60]. To test whether the reduced feedforward inhibition is mediated by cell loss we counted the number of PV^+^ interneurons in the CA2 region using double immunofluorescence staining with PV and RGS14. The results, however, showed no evidence for a change in PV^+^ interneuron density in the demyelinated hippocampus (**Supplementary Fig. 4**).

Together, the data suggest that feedforward inhibition is impaired while glutamatergic excitation is maintained, in part consistent with the C1q-tagged and microglia/macrophage stripped vGAT^+^ release sites (**Figs. 4,5**).

The cuprizone-induced switch from a net feedforward inhibition to excitation was prominently visible when stimulating SCs with a burst (5 stimuli @100 Hz, in physiological extracellular solution, **Fig. 6d**). In control CA2 PNs, the PSP peak amplitudes summated highly sublinear and the 5^th^ peak potential amplitude was on average ∼5 mV more hyperpolarized relative to the first peak potential due to slow inhibitory potentials [52] whereas after cuprizone treatment the PSP peak amplitudes summated and increased by ∼1 mV relative to the first peak (**Fig. 6d**). The strong feedforward inhibitory drive from CA3 typically limits CA2 spike output and in accordance APs were only observed in 2/14 control CA2 PNs at the maximum strength of 90 V (on average 0.5 ± 0.43 APs, **Fig. 6e**). A similar low probability for spike output of CA2 PNs was noted in cuprizone treated mice (1/10 recordings, on average 0.22 ± 0.22 APs, mixed-effect ANOVA, Treatment *F*_9,207_ = 0.297, *P* = 0.975, **Fig. 6e**). As expected GABA_A/B_ block, in control CA2 PNs a larger number of APs were evoked with SC stimulation (> 6 APs from 30 V, **Fig. 6e**). In striking contrast, however, following cuprizone treatment CA2 PNs showed in comparison to controls PNs a ∼3-fold lower spike output rate (**Fig. 6e**). Thus, despite the lower dynamic range in CA2 feedforward inhibition, the glutamate-mediated synaptic drive from CA3 neurons in the demyelinated hippocampus produces only a weak spike output.

### Reduced excitability of CA2 pyramidal neurons

To test whether the synaptic changes are accompanied by morphological changes of CA2 PNs we analyzed the current-clamp recordings and post-hoc reconstructed RGS14^+^ or PCP4^+^ neurons (**Fig. 7a**). The total RGS14^+^ area (stratum pyramidale and oriens) was not different following cuprizone treatment (**Supplementary Fig. 4**). Detailed reconstructions of biocytin-filled CA2 pyramidal neurons showed that the total dendritic length was not different (control, 4.28 ± 0.75 mm, *n* = 5 vs. cuprizone 3.28 ± 0.58 mm, *n* = 6, Mann-Whitney test U = 9, *P* = 0.329). However, Scholl analysis showed that cuprizone treatment was associated with a significantly redistribution of dendritic branches, with a lower number in the lacunosum moleculare, reflecting possibly dendritic atrophy and loss of input sites from the enthorinal cortex (*P* < 0.0001, **Fig. 7b**). When using depolarizing current injections in the soma, PNs showed the characteristic delay in spike generation typical as described for CA2 PNs previously [52]. The resting membrane potentials were not different (control, –81.4 ± 2.0 mV, *n* = 13 versus –82.6 ± 1.2, *n* = 11, unpaired t-test *P* > 0.60) but the maximum firing rate was lower in CA2 PNs from cuprizone-treated mice (**Fig. 7c**). The minimum current to evoke AP generation was, however, not different between groups (Mann-Whitney test U = 51, *P* = 0.565, *n* = 10 cuprizone and 12 control neurons, 5 mice/group). Interestingly, the sag ratio, as measured by the degree of depolarization upon hyperpolarization steps to –110 mV revealed that CA2 PNs from cuprizone-treated animals exhibited significantly increased depolarizing amplitudes, suggesting an increased dendritic hyperpolarization-activated cyclic nucleotide-gated (HCN) conductance (**Fig. 7d**). These changes were not associated with the expected reduction in neuronal input resistance (unpaired t-test *P* > 0.72, **Fig. 7e**). Finally, detailed analysis of single APs showed that most properties were similar except the first rate-of-rise component, reflecting the local axonal charging of the spike[61], which significantly decreased by on average ∼100 V s^−1^ in cuprizone-treated mice (**Fig. 7f**). Taken together, these data suggest that increased dendritic resting conductance, decreased dendritic surface area and reduced spike generation at the axonal output site limits spike output from CA2 pyramidal neurons.

**Figure 7.**
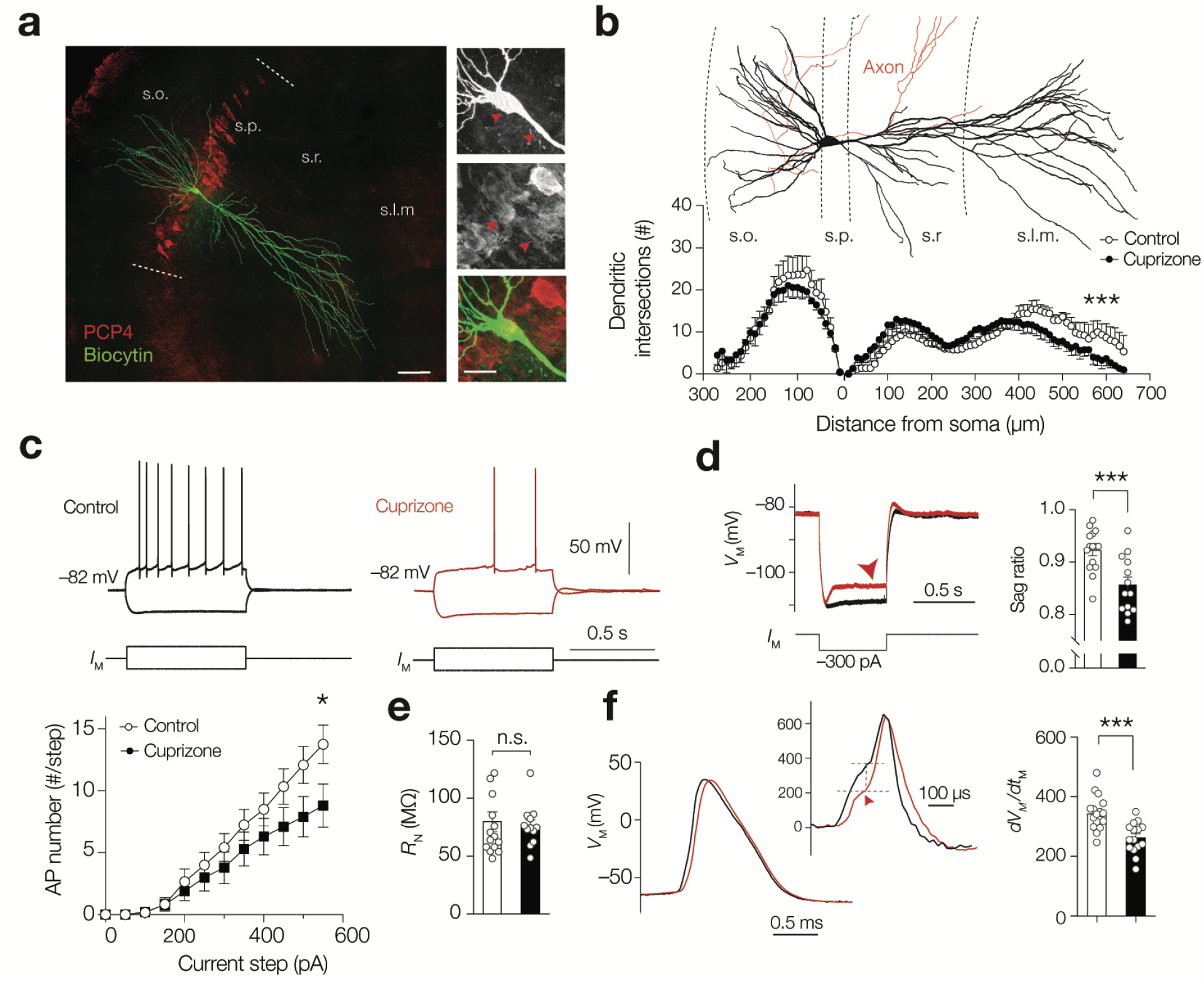
Cuprizone-induced demyelination reduces dendritic input sites and action potential generation in CA2 pyramidal neurons. **a**. Immunofluorescence image of a whole-cell recorded and biocytin-filled pyramidal neuron (streptavidin-biocytin, green), which was positive for the CA2 marker PCP4 (red). Scale bar, 100 µm. Inset scale, 25 µm. Red arrows indicate somatic and dendritic PCP4 expression. **b**. Top, example 3D morphological reconstruction of a CA2 pyramidal neuron from a control hippocampus. Axon in red. Bottom, Scholl plot analysis of control versus cuprizone neurons (*n* = 5 control neurons, *n* = 6 cuprizone), revealing a differential distribution of dendrites in the apical dendrites (mixed-effect RM ANOVA Treatment × Scholl ring interaction *F*_64, 530_ = 1.85, \*\*\**P* < 0.0001, Treatment *F*_1, 9_ = 0.088, *P* < 0.77) but not in basal dendrites (Treatment × Scholl interaction *F*_27, 222_ = 0.70, *P* = 0.863). **c**. Characteristic CA2 pyramidal neuron spike generation of a control CA2 and cuprizone-treated CA2 PN, showing delayed action potentials and near threshold ramp depolarization. Current-frequency (*I-f*) plots for CA2 PNs shows cuprizone reduces the maximum spike output rate (two-way RM ANOVA Treatment *P* = 0.1799, Treatment × current step interaction *P* = 0.0044, Holm-Sidak’s multiple comparison tests, for 0 to 500 pA *P* > 0.088, 550 pA \**P* = 0.022, cuprizone, *n* = 10 and *n* = 12 control neurons from 5 mice/group). Data represent mean ± SEM. **d**. Cuprizone increased the sag ratio to hyperpolarized steps near –110 mV (unpaired t-test, two-tailed *t*-test \*\*\**P* = 0.0007, *n* = 10 cuprizone and 15 control neurons, 5 mice/group). **e**. Input resistance (*R*_N_) was similar between groups (two-tailed *t*-test *P* = 0.898, *n* = 10 cuprizone and 15 control neurons, 5 mice/group). **f**.Action potentials are slower at their onset rate d*V*/d*t* (inset), reflecting axonal charging of the somatodendritic domain (two-tailed *t*-test \*\*\**P* = 0.0002, *n* = 15 cuprizone and 17 control neurons, 5 mice/group). For **d, e, f**, Bars represent mean ± SEM with circles individual cells.

### Social memory is impaired by cuprizone treatment

The dorsal CA2 area receives unique subcortical inputs signaling emotional states and is a circuit for social memory formation [32], mediated by suppressing the PV-mediated feedforward inhibition and gating CA2 output to the ventral CA1 area [32, 57, 62]. Previous studies showed that cuprizone treatment increases social behaviors in a resident-intruder paradigm, but negatively impacts on complex motor tasks and hippocampal-dependent spatial learning [45, 63]. To investigate directly whether the CA2-mediated encoding of social memory is affected we used the five-trial social memory test [32]. The results showed that cuprizone significantly affected social memory (two-way RM ANOVA *F*_1,20 =_ 13.7, *P* = 0.0014, *n* = 11 mice/group, **Fig. 8**). Whereas control mice significantly reduced the time investigating the familiar conspecific, indicating social memory, they dishabituated when confronted with a novel mouse (Bonferroni’s multiple comparison tests Trials 2–4, *P* < 0.034, **Fig. 8**, **<u>Supplementary Movie S1</u>)**. In contrast, cuprizone treatment for seven weeks caused a lack of habituation (Bonferroni’s multiple comparison tests for Trials 2–4 versus Trial 1, *P* > 0.60, *n* = 11, **Fig. 8**, **<u>Supplementary Movie S2</u>**), and neither did mice dishabituated with a novel mouse (*P* > 0.999, Trial 5 vs Trial 1, *n* = 11, **Fig. 8**).

**Figure 8.**
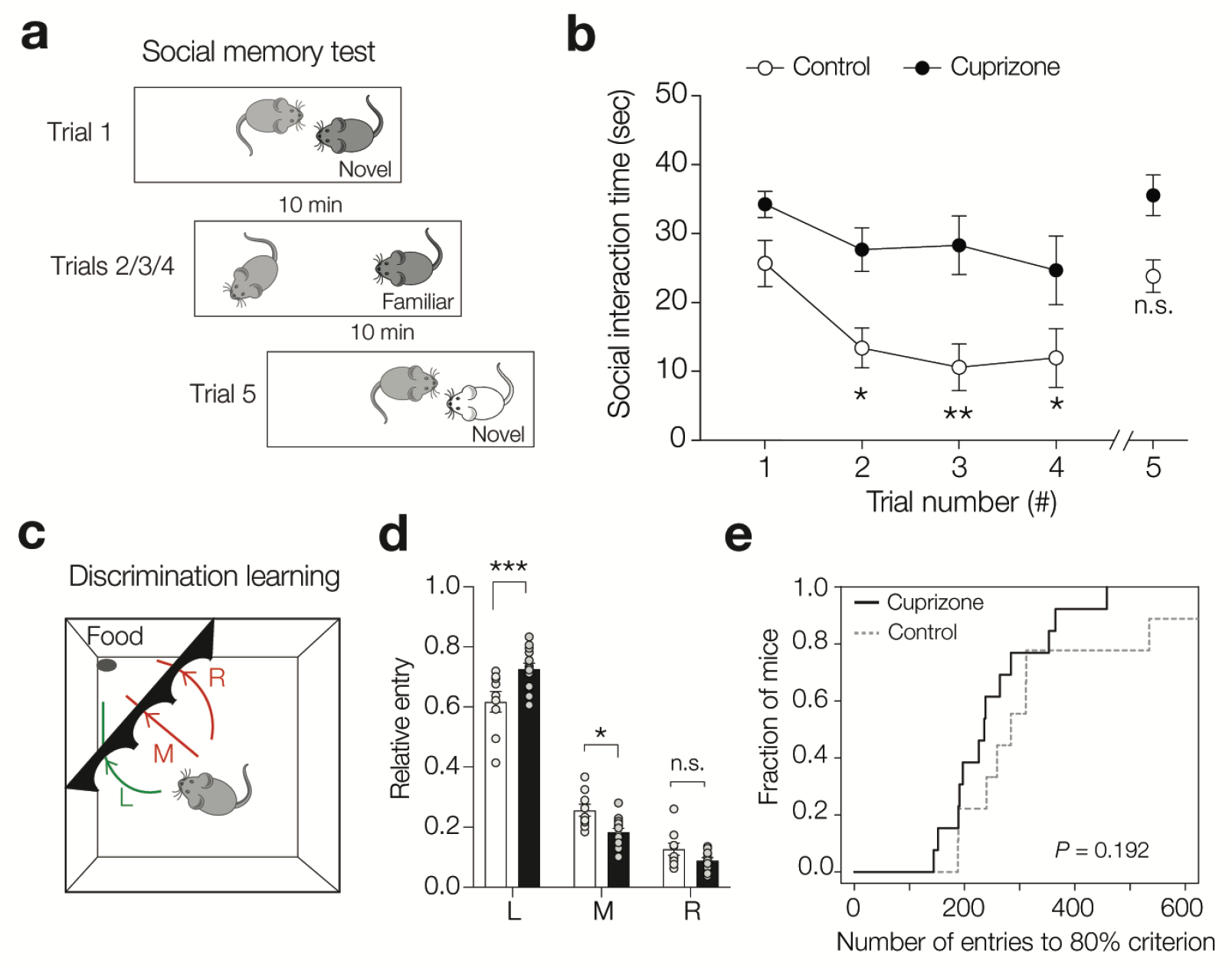
Cuprizone-induced demyelination impairs social memory but not discrimination learning. **a**. Schematic of the five-trial social memory test. Subject animals (light grey) were placed in a cage and after 15 min an unfamiliar novel mouse was introduced for a 1-minute trial duration. The stimulus reintroduced 4 times (trials 2 to 4) with each 10 minutes inter-trial intervals. Subject mice are expected to show habituation based on memorizing social cues. At the 5^th^ trial a novel mouse was introduced to measure dishabituation. **b**. Population analysis for the total time spent by the subject mouse socially investigating the stimulus mouse (anogenital sniffing, approaching and close proximity behaviors). Cuprizone treatment caused a significantly different habituation behavior (two-way repeated measures ANOVA, Treatment *F*_1,20_ =13.72, *P* = 0.0014, Trial × Treatment *F*_4,80 =_ 0.78, *P* = 0.779). Control mice habituated in trials 2, 3 an 4 (Bonferroni’s multiple comparisons tests compared to trial 1, *t* = 4.39, \**P* = 0.010; *t* = 4.91, \*\**P* = 0.0049; *t* = 3.69, \*\**P* = 0.034, respectively) and dishabituated in trial 5 (*t* = 0.66, *P* > 0.99, *df =* 10, *n* = 11 mice). In contrast, cuprizone mice did not show habituation compared to control trial 1 (Bonferroni’s multiple comparisons tests, *t* = 2.01, \**P* = 0.56; *t* = 1.16, *P* > 0.99; *t* = 1.83, *P* = 0.778 and *t* = 0.33, *P* > 0.99, respectively, *df =* 10, *n* = 11 mice). Data show mean ± SEM. **c**. Schematic design of the automated PhenoTyper arrangement showing the cognition wall with three ports and an automated food pellet dispenser. Mice needed to engage in a continuously running task to be rewarded with a pellet food, automatically dispensed when the mouse ran 5 times through the left port (green line, discrimination stimulus). **d**. Cuprizone-treated mice showed increased preference for the left and middle ports (ordinary two-way ANOVA Treatment × port interaction *F*_2, 60_ = 12.48, *P* < 0.0001. Data represent mean ± SEM with individual mice indicated with circles. **e**. Cumulative distribution plot of the fraction of mice versus total number of the total number of entries until 80% learning criterion (Chi-square test = 1.85, *P* = 0.192, *n* = 9 control and *n* = 13 cuprizone mice).

To test whether cuprizone-treated mice experience general deficits in motor activities or other learning tasks, not involving social memory, we used the PhenoTyper which for a continuous period of 16 hours automatically monitored mice for locomotion and spatial exploration. In addition, individually housed mice engaged in a cognition wall monitoring a discrimination learning task in which food pellets were provided as a reward [64]. Analysis of the general behavior showed that control and cuprizone-treated mice exhibited similar levels of exploration, were spending equal amounts of time outside the shelter box and moved similar distances in the cage across the 16 hours of recording (**Supplementary Fig. 4**). Interestingly, cuprizone-treated mice showed a higher preference for the left port, learning the food-reward discrimination more accurate and faster (*P* < 0.0001, **Fig. 8** and *P* < 0.0001, **Supplementary Fig. 5**, respectively).

Together, the findings indicate that cuprizone-induced demyelination causes deficits in learning in specific domains and, in line with the impaired CA2 inhibitory circuit, social cognition was found to be impaired.

## Discussion

In this study we identified the CA2 hippocampal subfield as a common target for C1q-associated loss of inhibitory synapses both in the MS hippocampus and cuprizone-induced demyelination. Using post-mortem MS tissue collected at rapid autopsy we found that C1q is most prominently increased in CA2, associated with a loss of vGAT^+^ synapses and its upregulation correlated with cognitive deficits. Although extremely valuable, the human MS post-mortem hippocampal tissue offers only an endpoint snapshot of a complex pathological cascade. Animal models which reproduce the synaptic alterations seen in MS are a prerequisite to elucidating the mechanisms and functional consequences of hippocampal changes. While acute EAE models reflect the contribution of acute inflammation to pathology [65], dietary cuprizone feeding in mice induces hippocampal demyelination with little inflammatory lesions [41, 42], resembling some of the histopathological presentations of grey matter lesions in MS patients [13]. Using both EM and confocal microscopy in the hippocampus of the cuprizone model of demyelination, we showed that C1q was selectively enriched at vGLUT2^+^ and vGAT^+^ synapses, which were engulfed by microglial processes resulting in a substantial reduction in the number of only GABAergic terminals in the stratum pyramidale and oriens of CA2. That GABAergic synapses are specifically tagged by C1q, engulfed and eliminated by microglia independently of anti-MOG antibodies in the cuprizone model, disconnects the process of C1q-mediated elimination of synapses from the classical role of C1q as recognition molecule of T-cell mediated antigen/antibody complexes that typically occurs in autoimmune diseases as part of the inflammatory response. This is in line with the current knowledge that MOG-antibody associated demyelinating disease is different from MS and cortical demyelination in MS occurs independent of anti-MOG antibodies [66]. Nonetheless, the association between the density of C1q protein localized in CA2 and the amount of MBP reduction in this region, in mice (**Supplementary Fig. 2c**) and in MS (**Fig. 7c**), may suggest that C1q-mediated processes are at least concomitant to myelin loss. Indeed, in the cuprizone model myelin debris itself suffices to activate Iba-1^+^ microglia/macrophages and causes hypertrophy of GFAP^+^ astrocytes in grey matter regions [67]. On the other hand, the process of microglia-mediated elimination of C1q-tagged synapses is reminiscent of observations during early development [18], adulthood [26], normal ageing [27] and neurodegeneration [22], all of which are conditions that do not involve demyelination, indicating that C1q-mediated elimination of synapses may also be an event independent of demyelination.

Recent work from Werneburg et al. [16] showed that synaptic material is tagged by complement C3 (not by C1q) and is engulfed by microglia in the retinogeniculate system of models of demyelination and in the visual thalamus of MS patients before onset of clinical disease and before overt signs of demyelination. In contrast to that study [16], we found that C1q but not C3 is deposited at discrete synapses that are engulfed by microglia in the MS and cuprizone hippocampus, suggesting that different complement pathways may be at play in different neural circuits in the mouse and human brain. Although we also observed a higher density of C3d in the CA2 hippocampal subfield of cuprizone mice compared to controls, this complement complex mainly localized at GFAP^+^ astrocytes. This is consistent with recent work showing that microglia-derived C1q (together with IL-1α and TNF) induce astrocytes to transition to a more reactive phenotype, including induction of astrocytic C3[51]. Interestingly, in the demyelinated CA2 region and within the stratum pyramidale and oriens layers, C1q specifically targets and eliminates vGLUT2^+^ and vGAT^+^ synapses but not Homer1^+^ or vGLUT1^+^ synapses. Whilst the vGLUT1 and Homer1 synapse markers were strongly increased across intrahippocampal regions (in cuprizone and MS hippocampi) our whole-cell recordings for Schaffer-collateral responses did not show a change in EPSP input strength. Interestingly, a recent study showed that Schaffer collateral field responses in CA1 reduce in amplitude in the first weeks of cuprizone feeding [68]. The discrepancies between functional recordings and glutamatergic synapse markers remain to be further investigated. One possibility is that the upregulation in these pre- and postsynaptic proteins are non-neuronal and represent reactive astrocytes. An upregulation of Homer1 also is seen in reactive astrocyte types that switches astroglial signalling pathways during inflammatory conditions [69].

An important question that arises from these findings is what molecular and/or activity-dependent mechanisms determine which synapses are targeted and which are spared by C1q? At the molecular level, a recent study in the retinogeniculate system demonstrated that a “don’t eat me” signal, such as CD47, is required to prevent excess pruning of synapses during development [70]. CD47 could directly inhibit phagocytosis by binding to its receptor, SIRPα, on microglia/macrophages [71, 72] and it has also been shown to prevent engulfment of cells opsonized with “eat me” flags, such as complement, showing that it can override these signals [73]. In addition, C1q was copurified with synaptosomes containing markers of apoptosis [74], suggesting that synaptic pruning may involve some of the same molecular triggers as the complement-mediated enhanced clearance of apoptotic cells that occurs as part of a homeostatic (non-phlogistic) process in the periphery [24]. Whether “don’t eat me” signals on spared synapses or “eat me” (i.e. apoptotic) signals on tagged synapses are involved in the engulfment of complement-tagged inhibitory synaptic elements in the rodent and/or MS demyelinated hippocampus remains to be determined. One additional mechanisms by which RGC synapses are eliminated during development involves neuronal activity, with microglia engulfing less active RGC inputs [19], in line with the knowledge that less active or ‘weaker’ inputs are pruned and lose territory as compared to those inputs that are ‘stronger’ or more active, which elaborate and strengthen [75]. The changes in ascending and descending inputs from the hippocampus as well as the intrahippocampal activity remains the be further examined.

### Functional reorganization of demyelinated intrahippocampal circuits

In the cuprizone model the loss of GABAergic terminals was not limited to specific CA subfields but more widespread. Indeed, cuprizone treatment affects besides the CA2-mediated social memory (this study, **Fig. 6**) also spatial navigation [45], typically mediated by synaptic plasticity in the CA1 and CA3 areas in the dorsal hippocampus in rodents[8]. Although CA2 represents only a small region of the CA pyramidal layer, emerging evidence points to CA2 as an hippocampal area that is genetically, molecularly and physiologically unique and acts as a hub controlling subcortical and intrahippocampal information processing[36], and could represent a novel target for therapeutic treatment [76]. Unlike CA1 and CA3, the CA2 PNs receive strong long-range extrahippocampal and input from both layer 2 medial and lateral entorhinal cortex pyramidal neurons at their distal dendrites [35, 52-54] and direct vGLUT2-mediated inputs from the medial septum diagonal band complex and the supramammillary nucleus [55, 56]. Whether the myelin loss from excitatory long-range projections cause impairments in the temporal structure of activity in CA2 needs to be determined by using *in vivo* recordings and cell-selective optogenetic approaches. Data on the electrophysiological consequences of myelin loss in the hippocampus is scarce. Recent longitudinal *in vivo* Ca^2+^ imaging of CA1 PNs over the course of 7 weeks cuprizone treatment reported a neuronal hypo-excitability, rapidly recovering during remyelination [68]. In contrast, using EEG recordings Hoffmann et al. [77], reported large-amplitude seizure activity in the hippocampus after 9 weeks of cuprizone treatment in awake and freely moving mice. The present finding of an impaired inhibitory circuit in the CA2 subregion (**Fig. 4**) could provide a cellular mechanism giving rise to hippocampal seizure activity. Consistent with this conjecture, Boehringer and colleagues [53] showed that the CA2 region acts as a central hub to balance excitation and inhibition across CA1 and CA3 areas. Using chemogenetic silencing of the CA2 neurons transformed sharp-wave ripple activity into seizure-like discharges [53]. Such widespread synchronization of inhibitory activity may in part be dependent on the strong feedforward inhibition by fast-spiking CA2 interneurons. The CA2 region contains a high density of PV^+^ interneurons [60, 78] with a unique morphology and mid-range axonal projections targeting CA1 and CA3 pyramidal layers. Interestingly, a large fraction of the intrahippocampal myelination represents sheaths that are wrapped around GABAergic interneuron axons, mostly including the PV^+^ axons [78, 79]. Whether PV^+^ interneuron excitability changes with demyelination and what triggers C1q upregulation to prune GABAergic release sites is not well understood but is an important area for further research.

### A role of CA2 circuits in cognitive impairments?

The importance of anatomical parcellation and functional subspecialization of the hippocampal subfields in cognitive problems in MS had already been brought forward by diffusion tensor MRI studies of atrophy and connectivity in MS [80]. For example, CA1 atrophy is a prediction for verbal memory performance [9, 10, 81], while CA2/3 atrophy underlies depressive symptoms [11], and changes in the dentate gyrus enlargement may explain poor cognitive performance[82]. The subcortical areas with which the CA2 pyramidal neurons and interneurons are connected with are involved in emotional regulation and include the hypothalamus, amygdala and septum. To what extent the CA2 connectivity and the local PV interneurons are affected in MS and whether its role in social memory consolidation is homologous to rodent is unknown and remains to be further examined with high-resolution MRI imaging enabling the parcellation of small CA2 area and by performing further detailed molecular analysis of area CA2 in the postmortem brain. The idea that CA2 in the human hippocampus is critical to cognition is, however, supported by a meta-analysis of post-mortem studies, revealing that PV^+^ interneuron loss specifically within the CA2 is one the strongest predictors for schizophrenia and mood disorders [83]. Both social cognition and facial emotion recognition, have been identified as domains that are affected in MS patients and associated with reduced social activities and a burden for the quality of life [5, 6]. Social cognition steers the ability to interpret and interact with the mental states of others and is a core psychological skill to maintain relationships and social support. This capacity is of crucial important for people with MS to mitigate the disease and recent studies showed that a decline in social cognitive skills negatively impacts on the quality of the life for MS patients. Our findings of a disrupted inhibitory CA2 circuit related to impaired memory for conspecifics in the demyelinated hippocampus resemble the changes in CA2 seen when deleting the psychiatric-disease related gene 22q11.2 causing also an impaired CA3 to CA2 feedforward mediated inhibition as well as a reduced CA2 PN spike output resulting in an impaired social memory [59]. Altogether, our work adds to the emerging evidence of subspecialization of the hippocampal subfields in specific cognitive domains which may also help explain subfield-specific susceptibility to injury in MS.

## Acknowledgments

We are grateful to the Electron Microscopy Centre Amsterdam (EMCA) of the Amsterdam UMC providing support with EM.

## Author Contributions

V.R. and M.K. jointly conceptualized the study; V.R., M.D., M.M., N.P., S.V. D. L., G.S. and M.K. developed the methodological approaches; V.R., J.L.G., J.J.G.G. and M.K. provided the lab resources, V.R., M.D., M.M., N.P., S.V., D.L., G.S. and M.K. conducted the experiments.; V.R., M.M., N.P., G.S. and M.K. analysed the data; V.R. and M.K. wrote the original draft; All authors contributed to reviewing and editing the manuscript, and approved its final version.

