## Supplementary information for "Complement-associated loss of CA2 inhibitory synapses in the demyelinated hippocampus impairs memory"

**Supplementary Figures**

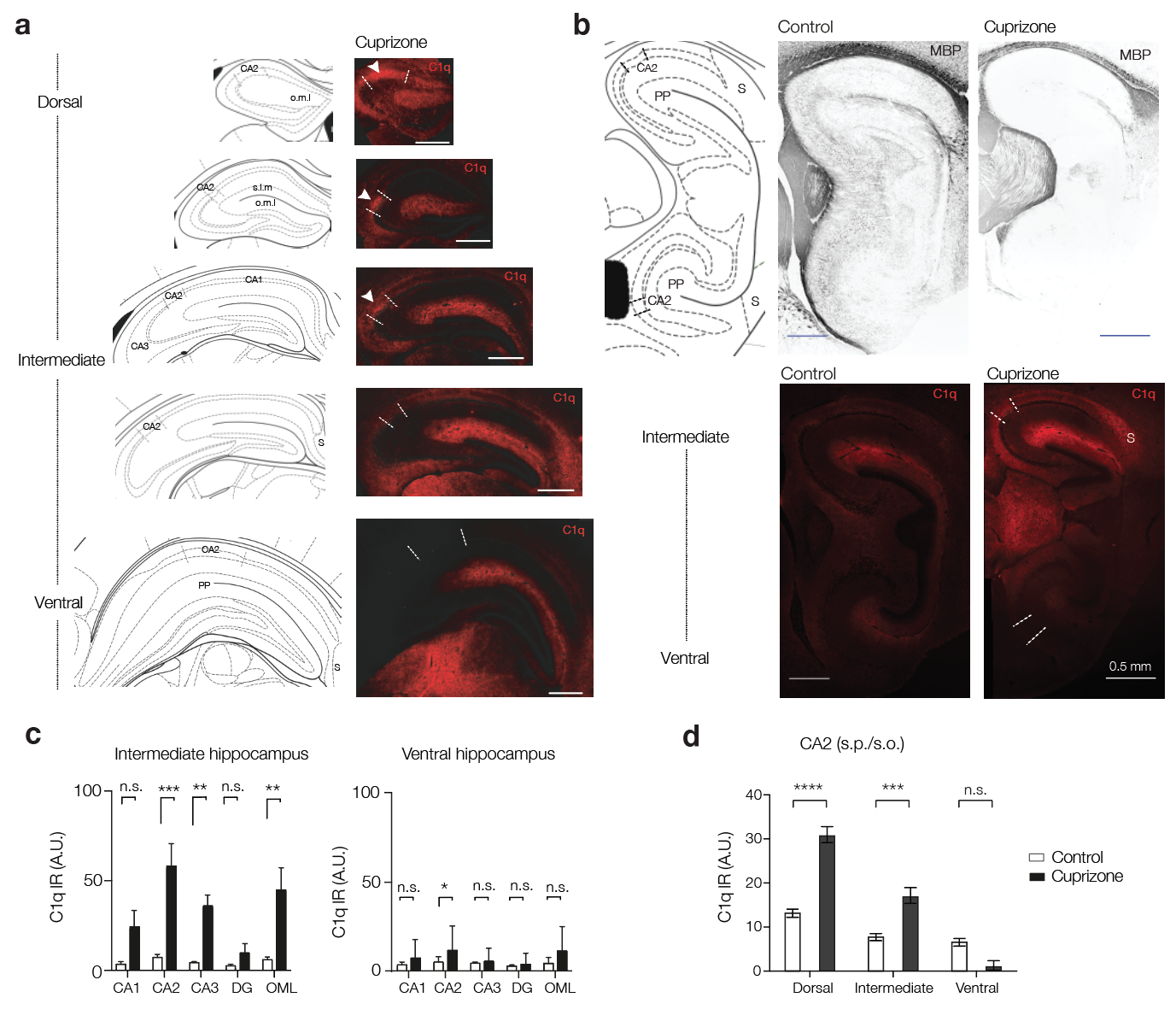

**Supplementary Figure 1. Dorsal to ventral gradient of cuprizone-induced C1q upregulation across the longitudinal axis of the hippocampus**

a. Left, coronal sections of the hippocampus adapted from the mouse brain atlas (Paxinos and Watson). Right, immunofluorescence images for C1q for the same locations shown on the left in cuprizone-treated mice (9 weeks, 0.2%). Note the high C1q intensity in the dorsal hippocampus (white arrows) but not in the ventral hippocampus (CA2 within dotted lines).
b. Sagittal view of the intermediate and ventral hippocampus. Note the difference between intermediate and ventral C1q intensity.
c. Population analysis of the hippocampal subfields for relative C1q intensity, as percentage area reveals a gradient in upregulation (two-way ANOVA with Sidak’s multiple comparison tests ****P* < 0.001, ***P* < 0.01, **P* < 0.05, n.s. = non significant, *n* = 6; 2 slices from 3 brains per group). Data shown as mean ± SEM. Closed bars, cuprizone and open bars control mice.
d. Comparative analysis of cuprizone-induced C1q upregulation across three anatomical levels of the CA2 subfield of the longitudinal axis of the hippocampus reveals a gradient in its increase (two-way ANOVA followed by Sidak’s post-hoc tests. Dorsal *****P* < 0.0001, intermediate ****P* = 0.0001, ventral *P* = 0.396, *n* = 6; 2 slices from 3 brains per group). C1q signals were averaged for the pyramidal and oriens layers. All data shows as mean ± SE

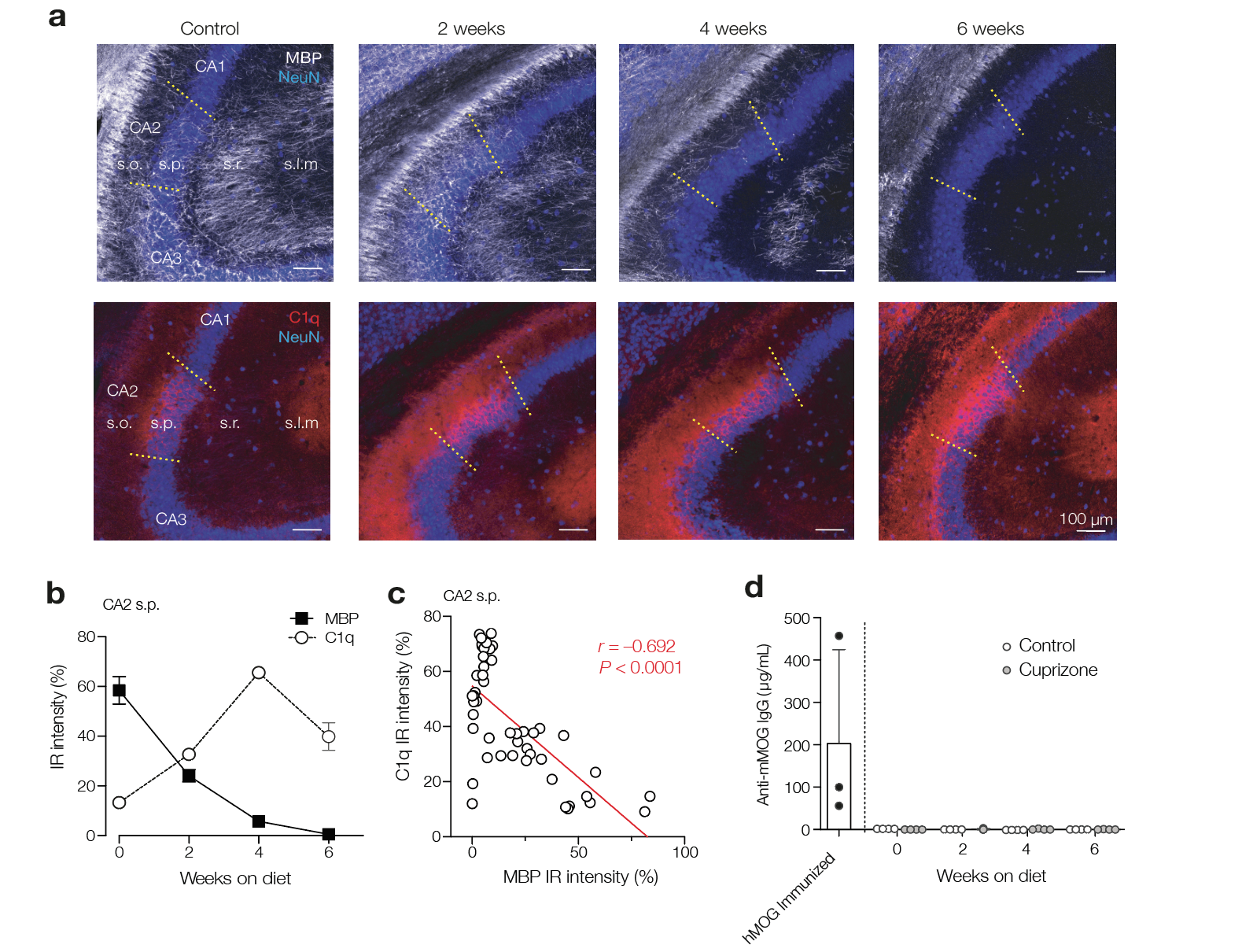

**Supplementary Figure 2. Temporal development of MBP and C1q in the CA2 subfield**

a. Example immunofluorescence images showing the dorsal CA2 hippocampal subfield for myelin basic protein (MBP, top, white) and C1q (bottom, red) overlaid with the neuronal marker NeuN (blue). The top and bottom row are the same brain sections from triple immunofluorescence staining separate for visual clarity. From left to right; control hippocampus and 2, 4 and 6-weeks treatment with the cuprizone diet.
b. Population data for the quantification of MBP and C1q intensities for the CA2 stratum pyramidale subfield in the dorsal hippocampus (48 images from 32 mice, 4 mice/group).
c. The cuprizone-induced MBP loss and C1q increase are negatively correlated across brain sections for the s.p. subfield (Pearson’s correlation coefficient *r* = –0.69, *n* = 48 sections from 4 mice per time/group). In contrast, neither in the stratum oriens (s.o.) nor in the stratum radiatium (s.r.) the myelin loss and C1q gain were significantly correlated (*P* = 0.160, slope = –0.350 and *P* = 0.102, slope = –0.0219, respectively, *n* = 48 sections from 4 mice/time/group).
d. ELISA was used to measure anti-mouse MOG IgG1 in the blood of control (*n* = 4) and cuprizone-fed (*n* = 4) mice throughout the period of cuprizone feeding up to 6 weeks. Sera from human myelin oligodendrocyte glycoprotein (MOG)-immunized C57BL/6 mice (*n* = 3) were collected during the chronic phase of experimental autoimmune encephalomyelitis (EAE) and used as an internal technical control

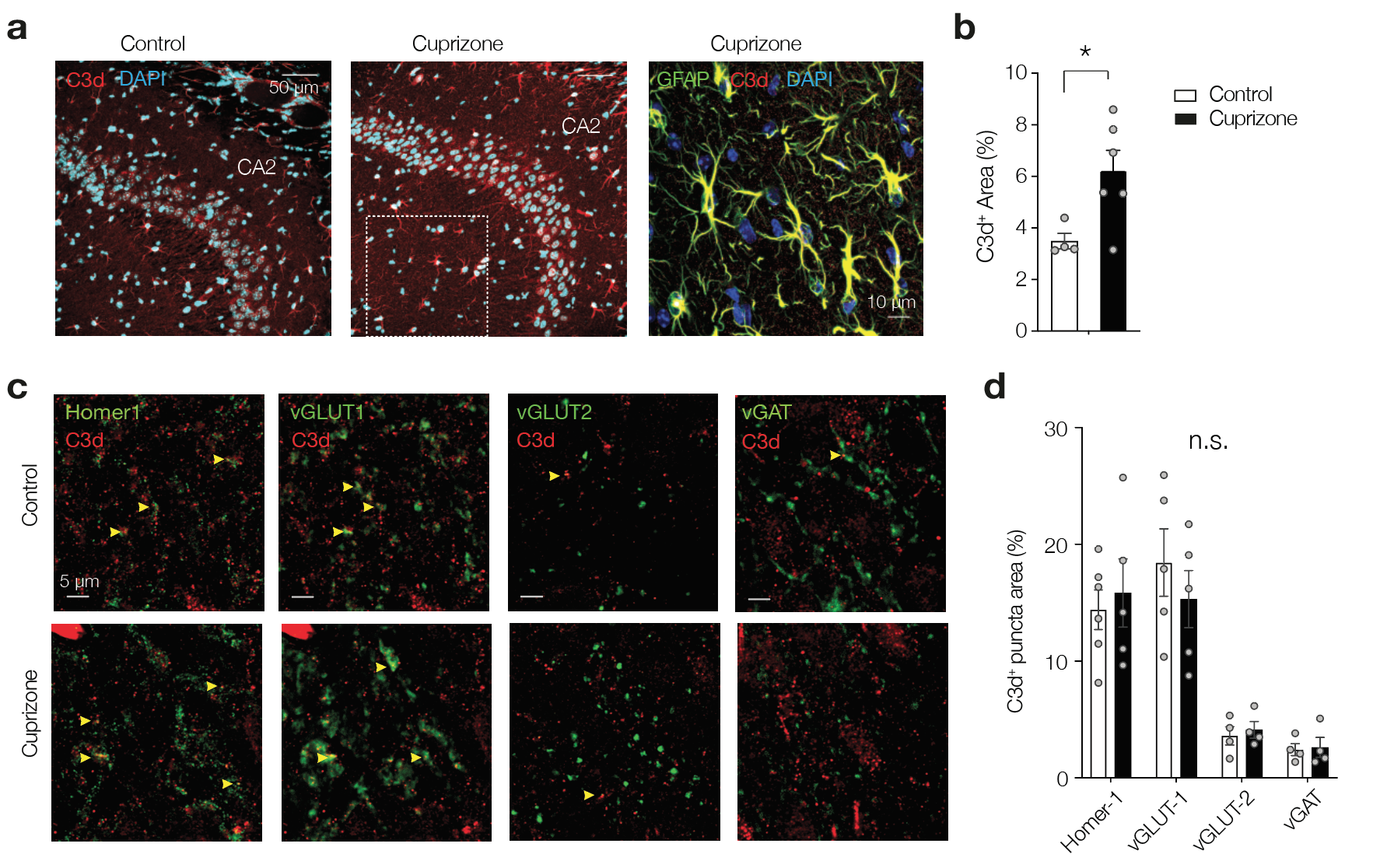

**Supplementary Figure 3. Demyelination-induced C3d upregulation preferentially targets vGLUT-1 containing glutamatergic synapses**

a. Left, double immunofluorescence staining for C3d (red) and DAPI (blue). Note that C3d is present in control hippocampus. Right, triple immunofluorescence staining for the astrocyte marker GFAP (green) reveals a strong overlap with C3d (red). Image is magnification from the inset in the left image (dotted square line).
b. Population analysis for the area covered by C3d in the CA2 subfield reveals a modest increase (unpaired *t*-test **P* = 0.03, *t* = 2.61, *df* = 8, *n* = 4 control and *n* = 6 cuprizone sections from 2 animals per group).
c. Cuprizone-induced synaptic changes were not associated with a differential change in C3d expression. Top row, control example immunofluorescent images of synaptic markers Homer1, vGLUT1, vGLUT2 and vGAT (green) co-stained with complement protein C3d (anti-C3d, red). Bottom row, similar staining combination and region of the CA2 area after 9 wks, 0.2% cuprizone. Arrowheads indicate overlap in expression (yellow). Scale bars, 5 µm.
d. Population analysis for area of co-localization. Note that C3d significantly clusters with vGLUT1 synapses (two-way ANOVA synapse type *F*_3,29_ = 25.50, *P* < 0.0001, Tukey’s multiple comparison tests vGLUT1 vs. vGLUT2 and vGAT, for both *P* < 0.0001, vGAT versus vGLUT2, *P* = 0.96). In contrast, complement complex C1q is more co-localized with vGLUT2^+^ and vGAT^+^ puncta (c.f. **Fig. 2d**). Overall, cuprizone-induced demyelination did not affect co-localization (two-way ANOVA treatment *F*_1,29_ = 0.028, *P* = 0.87) and neither showed interaction of treatment and synapse types (type × treatment *F*_3,29_ = 0.51, *P* = 0.67).

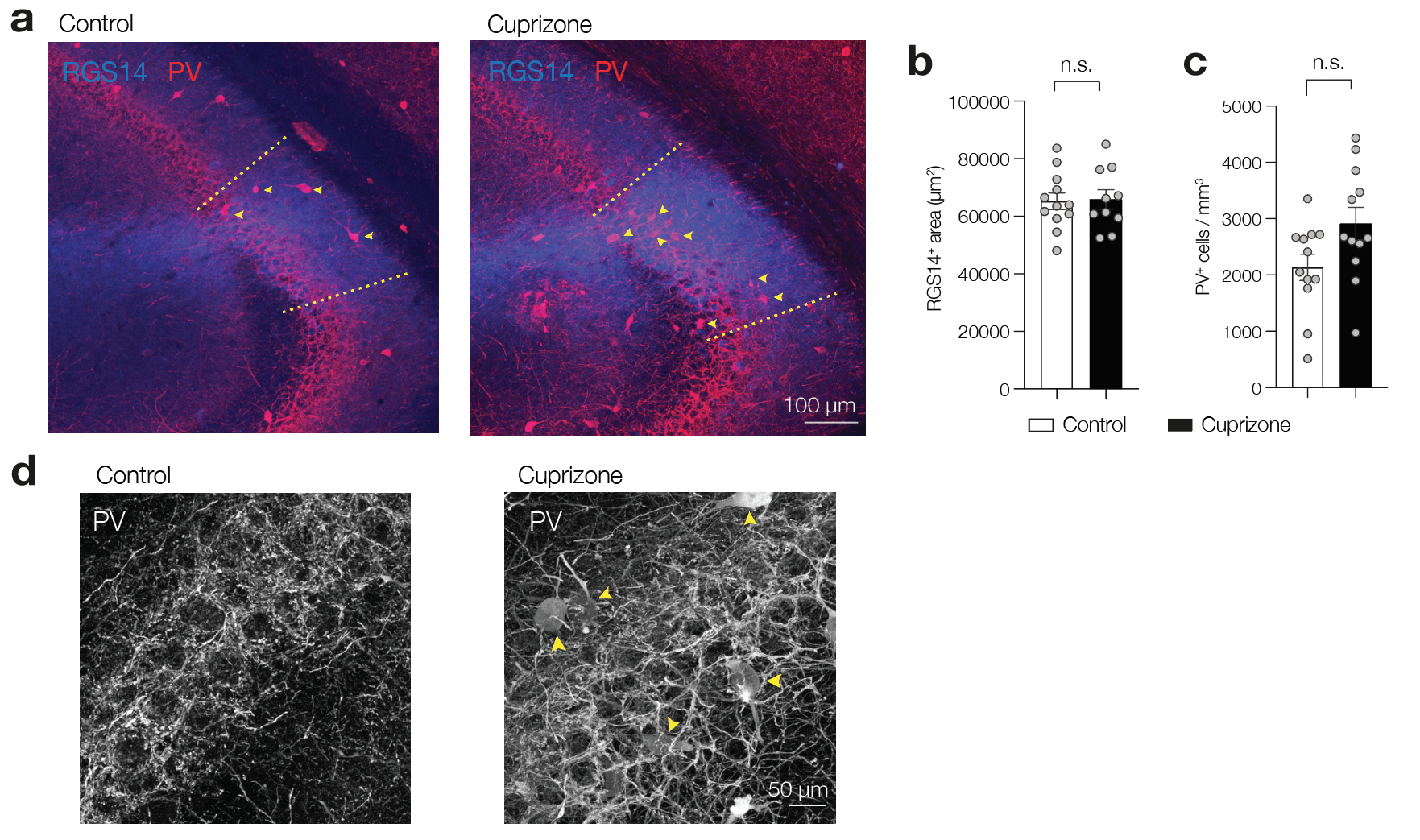

**Supplementary Figure 4. Cuprizone treatment does not impact the CA2 area or cell number**

a. Double immunofluorescence for RGS14 (blue) and parvalbumin (anti-PV, red) in control hippocampus (left) and the hippocampus after 7-weeks 0.2% cuprizone treatment (right). The borders of the CA2 region are indicated with dashed yellow line. PV^+^ somata included in the analysis are indicated with yellow arrows.
b. Population analysis for the RGS14^+^ area as an indication for the CA2 field size shows no difference (two-tailed Mann-Whitney test U = 56, *P* = 0.377, *n* = 12 sections for 6 mice/group. For each hemisphere 10 consecutive sections of 40 µm in the left and right dorsal hippocampus were stained and averaged.
c. Population analysis for the PV^+^ neurons in CA2 reveals no change with cuprizone-induced demyelination (two-tailed Mann-Whitney test U = 43, *P* = 0.101, *n* = 12 sections for 6 mice/group). Mean ± SEM. Circles show individual hippocampus, left and right.
d. Higher magnification of the CA2 region and PV immunofluorescence (grey). Yellow markers indicate PV^+^ cell bodies.

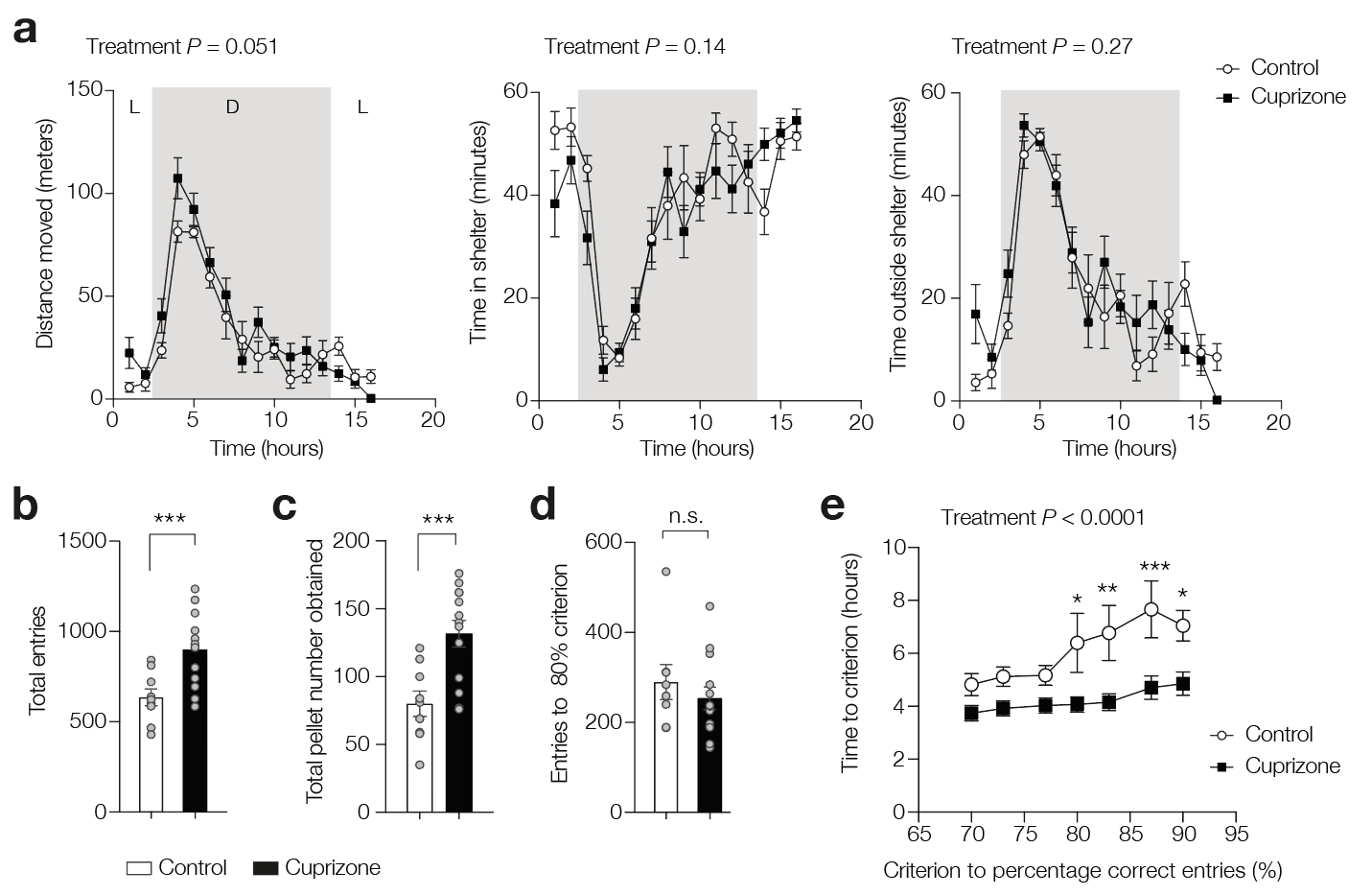

**Supplementary Figure 5. Cuprizone treatment does not affect general activity while improving spatial discrimination learning**

a. One-hour binned plots of the parameters (from left to right) distance moved, time inside shelter and time outside the shelter box over 16 hours continuous recording of mice individually housed inside the PhenoTyper cage. Cuprizone treatment did not change activity patterns (distance moved; two-way ANOVA *F*_1, 20_ = 4.33, Treatment *P* = 0.051, Time × treatment *F*_15, 300_ = 1.71, *P* = 0.048). Time within the shelter; mixed-effect model *F*_1, 319_ = 2.14, *P* > 0.145 and time outside shelter; mixed-effect model *F*_1, 319_ = 1.093, *P* = 0.296, respectively, *n* = 9 control and *n* = 13 cuprizone mice). Data and connecting lines show mean ± SEM. Note that both groups follow similar patterns across light (L) and dark (D) phases of the circadian cycle.
b. Cognition wall behavior for the control (open bars) and the cuprizone-treated mice (closed bars). Over the recording period cuprizone-treated mice made more movements through the entrances of the cognition wall (Two-tailed unpaired *t*-test with Welch’s correction (*t, df*) = 3.6, 20.0, *P* = 0.0018)
c. Cuprizone-treated mice earned ~60% more pellets (two-tailed unpaired *t*-test with Welch’s correction (*t, df*) = 3.8, 19.6, *P* = 0.0011).
d. The total number of entries through all three ports (L, M and R) until a 80% criterion was observed was not different (two-tailed unpaired *t*-test with Welch’s correction (*t, df*) = 0.775, 12.83; *P* = 0.451). Learning criteria was assessed by a moving average over the last 30 entries in which a specific percentage (80%) of entries through the left (L) entry port (providing the food reward) was achieved.
e. Cuprizone-treated mice more rapidly learned to discriminate the cognition wall task. Time until a specific cut-off learning criterium was reached plotted against the learning criterion between 70 and 90% correct entries. Ordinary two-way ANOVA Treatment *F*_1, 132_ = 46.29, ****P* < 0.0001, Percentage *F*_6, 132_ = 3.85, ***P* = 0.0014, Interaction *F*_6, 132_ = 1.088, *P* = 0.37. Note the cuprizone treated mice reach the learning cut-off before control mice. Sidak’s multiple comparison tests for < 77% *P* > 0.6, for 80% **P* = 0.0157, 83% ***P* = 0.0043, 87% ****P* = 0.0008, 90% **P* < 0.0382 (for all, *n* = 8 control, *n* = 13 cuprizone mice)

**Supplementary movies**

**Supplementary movie S1**

Example movie of a control mouse in the third trial within a five-trial social test. A familiar mouse is placed in the cage for the duration of 1 minute.

**Supplementary movie S2**

Example movie of a cuprizone-treated mouse in the third trial. The familiar mouse is placed in the cage for the duration of 1 minute. Note the long duration of anogenital sniffing and frequent approaching.

| **SUPPLEMENTARY TABLES**  **Supplementary Table 1. Clinical data of MS donors and controls** | | | | | | | |
| --- | --- | --- | --- | --- | --- | --- | --- |
| **Case** | **Sex**  M/F | **Age**  (years) | **PMD**  (hours) | **DD**  (years) | **MS type** | **Cognitive decline** | **COD** |
| MS |  |  |  |  |  |  |  |
| *Myelinated* |  |  |  |  |  |  |  |
| 1993-307 | F | 72 | 9.50 | 41 | PP | Unknown | Baroreflex insufficiency |
| 1995-276 | M | 56 | 5.75 | 13 | SP | Unknown | Respiratory insufficiency |
| 1996-264 | F | 67 | 4.90 | 2 | PP/SP | Unknown | Euthanasia |
| 1996-352 | F | 53 | 7.25 | 18 | SP | Unknown | Pneumonia |
| 1998-087 | F | 55 | 7.90 | 11 | PP/SP | Unknown | Aspiration pneumonia |
| 1998-185 | F | 70 | 8.90 | 19 | PP | Unknown | Cardiogenic shock & pneumonia |
| 1999-025 | F | 64 | 7.75 | 35 | SP | Unknown | Dehydration & pneumonia |
| 2000-124 | M | 64 | 7.50 | 34 | PP | Unknown | End stage MS |
| 2002-025 | M | 77 | 4.25 | 26 | SP | Unknown | Cerebral vascular accident |
| 2002-053 | F | 48 | 5.85 | 21 | PP/SP | Unknown | Congestive cardiac failure |
| 2003-106 | F | 72 | 8.85 | 20 | SP | Unknown | General deterioration |
| 2004-055 | M | 71 | 7.00 | 26 | PP/SP | Unknown | Pneumonia by aspiration |
| 2007-001’ | F | 57 | 23.05 | 21 | PP/SP | + | Sepsis |
| 2007-002’ | F | 78 | 11.10 | 30 | SP | - | Dehydration |
| 2007-071’ | M | 41 | 7.20 | 14 | PP | + | Urosepsis and pneumonia |
| 2009-036’ | M | 72 | 7.55 | 23 | SP | - | Pneumonia |
| 2009-103’ | F | 59 | 4.45 | 24 | SP | - | Euthanasia |
| 2010-024’ | M | 44 | 10.15 | 22 | SP | + | General deterioration |
| 2011-120’ | M | 73 | 8.45 | 51 | SP | + | Urosepsis |
| 2012-032’ | M | 80 | 9.45 | 36 | SP | + | Cachexia and pneumonia |
| 2013-047’ | F | 48 | 11.50 | 22 | SP | + | Respiratory failure |
| 2013-081* | M | 66 | 10.55 | 25 | PP | Unknown | Euthanasia |
| 2013-032’ | F | 87 | 9.30 | 20 | PP/SP | - | Dehydration and Renal insufficiency |
| 2014-052* | F | 61 | 4.15 | Unknown | PP/SP | Unknown | Euthanasia |
| 2014-057* | M | 61 | 5.15 | Unknown | PP/SP | Unknown | Euthanasia |
| 2015-020* | F | 71 | 3.52 | Unknown | PP/SP | Unknown | Euthanasia |
| 2015-022* | M | 83 | 3.00 | Unknown | PP/SP | Unknown | Pneumonia |
| 2015-064* | M | 51 | 5.30 | Unknown | PP/SP | Unknown | Pneumonia |
| 2015-070* | F | 77 | 4.00 | Unknown | PP/SP | Unknown | Pneumonia |
| 2016-017* | M | 61 | 5.09 | Unknown | PP/SP | Unknown | Euthanasia |
| 2017-083* | F | 81 | 2.30 | Unknown | PP/SP | Unknown | Anorexia |
| *Demyelinated* |  |  |  |  |  |  |  |
| 1992-187 | F | 35 | 5.75 | 11 | SP | Unknown | General decline |
| 1996-116 | F | 81 | 4.25 | 49 | SP | Unknown | Cachexia |
| 1996-076 | M | 46 | 3.75 | 23 | SP | Unknown | Pneumonia |
| 1997-123 | F | 89 | 5.90 | 12 | PP/SP | Unknown | Respiratory tract infection |
| 1998-059 | F | 76 | 4.60 | 2 | PP/SP | Unknown | Uremia; dehydration |
| 1999-086 | M | 81 | 8.85 | 59 | PP | Unknown | General deterioration |
| 2001-001 | F | 68 | 7.85 | 16 | PP/SP | Unknown | Pneumonia |
| 2001-137 | F | 56 | 6.90 | 9 | PP | Unknown | Urosepsis |
| 2003-105 | M | 66 | 7.50 | 26 | PP | Unknown | Unknown |
| 2004-017 | M | 49 | 8.00 | 25 | SP | Unknown | Pneumonia by MS |
| 2004-035 | M | 47 | 7.25 | 7 | SP | Unknown | Urosepsis with organ failure |
| 2008-053’ | F | 64 | 10.10 | 39 | SP | - | Urinary tract infection |
| 2008-096’ | F | 88 | 7.55 | 34 | PP | - | Exhaustion by chronic colitis ulcerosis |
| 2009-007’ | F | 67 | 9.15 | 42 | SP | - | Euthanasia |
| 2011-080’ | F | 56 | 8.25 | 34 | PP | + | Pneumonia |
| 2015-006* | F | 53 | 4.00 | Unknown | PP/SP | Unknown | Euthanasia |
| 2015-108* | F | 82 | 3.40 | Unknown | PP/SP | Unknown | Euthanasia |
| 2016-050* | F | 52 | 3.40 | Unknown | PP/SP | Unknown | Euthanasia |
| 2016-052* | F | 75 | 5.15 | Unknown | PP/SP | Unknown | Respiratory failure |
| 2016-074* |  | 65 | 4.15 | Unknown | PP/SP | Unknown | Stroke |
| 2017-001* | F | 40 | 2.30 | Unknown | PP/SP | Unknown | Euthanasia |
| 2017-068* | F | 49 | 3.55 | Unknown | PP/SP | Unknown | Pneumonia |
| 2017-139* | M | 75 | 4.00 | Unknown | PP/SP | Unknown | Euthanasia |
| 2017-161* | F | 65 | 3.50 | Unknown | PP/SP | Unknown | Fever |
| Controls |  |  |  |  |  |  |  |
| *Non-neurological* |  |  |  |  |  |  |  |
| 1997-115 | M | 81 | 7.90 | - | - | - | Renal insufficiency & heart failure |
| 1998-003 | F | 80 | 7.00 | - | - | - | Pulmonary embolisms |
| 2005-068 | F | 50 | 4.15 | - | - | - | Metastasized cell bronchocarcinoma |
| 2008-027 | F | 50 | 6.85 | - | - | - | Metastasized breast carcinoma |
| 2009-095 | F | 61 | 8.80 | - | - | - | Euthanasia |
| PMD, Post-mortem delay; DD, Disease duration; MS, Multiple Sclerosis; COD, Cause of death; PP, Primary progressive; SP, Secondary progressive; F, Female; M, Male; AD, Alzheimer’s Disease. The asterisk * indicates cases used for pathological/MRI correlation studies. The apostrophe’ indicates cases used for pathological/cognitive correlation studies. | | | | | | | |

| **Supplementary Table 2. Primary antibodies, dilution, source for immunohistochemistry** | | | | |
| --- | --- | --- | --- | --- |
| Antigen | Clone | Dilution | Source | Identifier |
| ***Mouse*** |  |  |  |  |
| Complement complex 1 (C1q) | Monoclonal 4.8 | 1:000 | Abcam | Cat# ab182451  RRID: AB_2732849 |
| NeuN | A60 | 1:1000 | Millipore | Cat# ABN90P  RRID: AB_2341095 |
| DAPI |  | 1.5 µg/ml | Vector | Cat# 7F0515 |
| Myelin Basic Protein (MBP) | 12 | 1:250 | Millipore | Cat# MAB386  RRID: AB_94975 |
| vGAT1 | Polyclonal | 1:500 | Millipore | Cat# AB5062P  RRID: AB_2301998 |
| vGLUT1 | Polyclonal | 1:1000 | Synaptic Systems GmbH | Cat# 135304  RRID: AB_887878 |
| vGLUT2 | 8G9.2 | 1:200 | Millipore | Cat# MAB5504  RRID: AB_2187552 |
| Homer1 | Polyclonal | 1:200  1:500 | Synaptic Systems GmbH | Cat# 160006  RRID: AB_2631222 |
| RGS14 | N133/21 | 1:500 | Neuromab | Cat# 75-170  RRID: AB_2179931 |
| Purkinje cell protein 4 (PCP4) | Polyclonal | 1:250 | Sigma | Cat# HPA005792  RRID: AB_1855086 |
| Streptavidin, Alexa Fluor 488^TM^ Conjugate |  | 1:400 | Invitrogen | Cat# S32354  RRID: AB_2315383 |
| C3d Complement | Polyclonal | 1:500 | Agilent | Cat# A006302  RRID: AB_578478 |
| Iba-1 | EPR16589 | 1:500 | Abcam | Cat# ab178847  RRID: AB_2832244 |
| GFAP | Polyclonal | 1:100 | Bio-connect | Cat# SC-6170  RRID: AB_641021 |
| ***Human*** |  |  |  |  |
| Proteolipid protein (PLP) | plpc1 | 0.3 μg/ml^a^ | Serotec | Cat# MCA839G  RRID: AB_2237198 |
| Human Leukocyte Antigen (HLA) | CR3/43 | 8.6 µg/ml^a^ | Abcam | Cat# Ab55152  RRID: AB_944199 |
| Vesicular glutamate transporter 1 (vGLUT1) | Polyclonal | 1:400^b^ | Synaptic  Systems | Cat# 135 304  RRID: AB_887878 |
| Vesicular GABA transporter 1 (vGAT) | Polyclonal | 1:100^b^ | Synaptic  Systems | Cat# 131 006  RRID: AB_2619820 |
| Post Synaptic Domain (PSD) 95 | D27E11 | 1:150^b^ | Cell  Signalling | Cat# 3450  RRID: AB_2292883 |
| Gephyrin | mAb7a | 1:100^b^ | Synaptic  Systems | Cat# 147 021  RRID: AB_2232546 |
| C1q | Monoclonal [34E2] | 1:100^b^ | Abcam | Cat # ab235454 |
| Antigen retrieval of paraffin sections was performed by heat in ^a^ 0.05 M Tris buffered saline pH 7.6 or ^b^ 10 mM Tris/1 mM EDTA buffer pH 9. | | | | |
